# Quantitative effects of environmental variation on stomatal anatomy and gas exchange in a grass model

**DOI:** 10.1101/2021.11.10.468085

**Authors:** Tiago DG Nunes, Magdalena W Slawinska, Heike Lindner, Michael T Raissig

## Abstract

Stomata are cellular pores on the leaf epidermis that allow plants to regulate carbon assimilation and water loss. Stomata integrate environmental signals to regulate pore apertures and optimize gas exchange to fluctuating conditions. Here, we quantified intraspecific plasticity of stomatal gas exchange and anatomy in response to seasonal variation in *Brachypodium distachyon*. Over the course of two years we (i) used infrared gas analysis to assess light response kinetics of 120 Bd21-3 wild-type individuals in an environmentally fluctuating greenhouse and (ii) microscopically determined the seasonal variability of stomatal anatomy in a subset of these plants. We observed systemic environmental effects on gas exchange measurements and remarkable intraspecific plasticity of stomatal anatomical traits. To reliably link anatomical variation to gas exchange, we adjusted anatomical *g*_s_max calculations for grass stomatal morphology. We propose that systemic effects and variability in stomatal anatomy should be accounted for in long-term gas exchange studies.

---

Stomata are the cellular pores on the leaf epidermis that allow plants to balance photosynthetic carbon dioxide (CO_2_) uptake with water vapor loss. Stomatal movements result from changes in turgor of stomatal cells (Jezek & Blatt, 2017). Stomatal opening is induced by an increase of turgor pressure in guard cells (GCs), while a decrease of turgor pressure in GCs results in stomatal closure. To optimize gas exchange, stomata interpret and integrate a plethora of environmental cues such as light, humidity, temperature, carbon dioxide concentration and even biotic factors like pathogens (Engineer et al., 2016; Jezek & Blatt, 2017; Kollist et al., 2014; Merilo et al., 2014; Murata et al., 2015; Sierla et al., 2016). In high light, for example, stomata of C_3_ and C_4_ plants open to provide sufficient CO_2_ for photosynthesis. In low light, on the other hand, less CO_2_ is required to saturate photosynthesis and, consequently, stomata close to limit water loss. Therefore, stomatal responsiveness and fast opening and closing kinetics can significantly contribute to plant water use efficiency (WUE) in changing environments (Lawson & Vialet-Chabrand, 2019; McAusland et al., 2016). WUE represents the ratio of carbon assimilation and water loss and is a crucial trait for plant productivity and stress resilience (Leakey et al., 2019; McAusland et al., 2016). Grasses, which include the cereals like rice, maize and wheat, show comparatively fast stomatal movements that likely contribute to more water-efficient gas exchange in changing environments (Franks & Farquhar, 2007; Lawson & Matthews, 2020; McAusland et al., 2016).

During the day plants face changing environmental conditions such as fluctuating ambient light intensity (𝒬_out_), temperature (T) and relative humidity (RH). Stomata mostly respond locally to environmental stimuli. This allows infrared gas analyser (IRGA)-based leaf gas exchange studies to be robust since leaves are placed in a chamber and exposed to controlled 𝒬_out_, RH, T and CO_2_ concentration ([CO_2_]) regardless of the ambient conditions. Nevertheless, it has already been suggested that external ambient conditions might systemically affect local stomatal responses measured by IRGA systems (Devireddy et al., 2018, 2020; Ehonen et al., 2020). This might be particularly relevant for gas exchange studies that are performed in greenhouse or field settings with significant daily and seasonal environmental fluctuations. However, the putative systemic influence of the varying ambient conditions on gas exchange measurements is not generally accounted for.

Furthermore, gas exchange parameters such as carbon assimilation (*A*), stomatal conductance to water vapor (*g*_sw_), intrinsic water use efficiency (iWUE) and stomatal kinetics are influenced by anatomical traits such as stomatal density (SD) and stomatal length (SL) (Elliott-Kingston et al., 2016; Faralli et al., 2019; Haworth et al., 2021; Lawson & Blatt, 2014). SD and SL are negatively correlated and vary in response to a variety of environmental conditions such as T, RH, [CO_2_] or 𝒬_out_ (Bertolino et al., 2019; Franks et al., 2009; L. Zhang et al., 2021). The seasonal variation of environmental conditions might, therefore, affect the intraspecific plasticity of stomatal anatomical traits influencing gas exchange performance and, consequently, the results of long-term gas exchange phenotyping studies.

Here, we quantified stomatal conductance kinetics in 120 individuals of the grass model *Brachypodium distachyon* (Bd21-3) in a greenhouse over the course of two years. Simultaneously, we logged the environmental conditions in the greenhouse (𝒬_out_, T, RH) and time of the day (time) and quantified how these parameters affected the measured gas exchange parameters (*A, g*_sw_, iWUE and stomatal response kinetics). We additionally quantified anatomical traits of stomata (SD and SL) in three different seasons (summer, autumn and winter) and observed a significant impact of seasonal growth conditions on these traits. This allowed us to correlate how variations in SD and SL influence steady-state gas exchange, stomatal kinetics and maximum stomatal conductance (*g*_s_max). When calculating anatomical *g*_s_max based on anatomical traits, we realized that existing approaches to calculate maximum pore area for the double end-correction version of the equation by Franks & Farquhar (2001) did not sufficiently account for the graminoid morphology. Using quantitative morphometry of open and closed *B. distachyon* stomata we determined how to accurately calculate maximum pore area and pore depth. These adjustments allowed for an accurate prediction of physiological *g*_s_max based on stomatal anatomical traits in *B. distachyon*.

## Material and methods

### Plant material and growth conditions

*B. distachyon* Bd21-3 seeds were vernalized in water for 2 days at 4°C before being transferred to soil. Plants were grown in a greenhouse with 18 h light:6 h dark, average day temperature = 28°C, average night temperature = 25°C and average relative humidity = 40 %. We used 6 cm X 6 cm X 8 cm pots per plant filled with 4 parts soil (Einheitserde CL ED73) and 1 part vermiculite. The greenhouse is located at 49° 24’ 52.38’’ N and 8° 40’ 5.808’’ E at the Centre for Organismal Studies Heidelberg, Im Neuenheimer Feld 360, 69120 Heidelberg, Germany. Daily mean temperature (January 2019-September 2021) varied between 3-5°C in December-February, 8-11°C in March-April, 13-21°C in May-September and 6-12°C in October-November (Deutscher Wetterdienst, https://cdc.dwd.de/portal/). Average daylight hours are 8.3-10.2 h in December-February, 11.9-13.8 h in March-April, 12.6-16.2 h in May-September and 9.1-10-8 h in October-November (Deutscher Wetterdienst, https://cdc.dwd.de/portal/).

### Leaf-level gas exchange measurements

All measurements were performed on *B. distachyon* leaves 3 weeks after sowing using a LI-6800 Portable Photosynthesis System (Li-COR Biosciences Inc.) equipped with a Multiphase Flash Fluorometer (6800-01A) chamber. The youngest, fully expanded leaf was measured using the 2 cm^2^ leaf chamber. Conditions in the LI-6800 chamber for light-response experiments were as follows: flow rate, 500 μmol s^−1^; fan speed, 10000 rpm; leaf temperature, 28°C; relative humidity (RH), 40 %; [CO_2_], 400 μmol mol^−1^; photosynthetic active radiation (PAR), 1000 – 100 – 1000 – 0 μmol PAR m^−2^ s^−1^(20 min per light step) (Fig. 1A). Light-response measurements of *A* and *g*_sw_ were obtained for 120 wild-type Bd21-3 individuals between May 2019 and September 2021. Gas exchange measurements were automatically logged every minute. Relative *g*_sw_ was calculated by normalizing *g*_sw_ to the highest *g*_sw_ value observed to evaluate kinetics of stomatal response regardless of variation on absolute *g*_sw_, eliminating the influence of stomatal density and leaf area. Because *B. distachyon* leaves do not fill the 2 cm^2^ chamber, individual leaf area was measured for a subset of 35 individuals to accurately quantify absolute *g*_sw_ and *A*. To obtain a mean approximation of gas exchange levels for the total 120 individuals, absolute *g*_sw_ and *A* were corrected by using the average leaf area (0.64 cm^2^) from the data subset (n=35). Intrinsic WUE (iWUE) was calculated as the *A* to *g*_sw_ ratio (*A*/*g*_sw_). Ambient light intensity (𝒬_out_) was monitored during the measurements using an external LI-190R PAR Sensor attached to LI-6800. Greenhouse temperature and relative humidity were monitored during the experiments using an Onset HOBO U12-O12 4-channel data logger that was placed next to the plants used for analysis. One-phase decay or one phase association non-linear regressions were obtained for the stomatal closure transitions (1000 – 100 and 1000 – 0 PAR) and stomatal opening transition (100 – 1000 PAR), respectively, to determine half-time (T_50%_) and rate constant (k). Maximum stomatal conductance (*g*_s_max) measurements were performed with the following conditions: flow rate, 500 μmol s^−1^; fan speed, 10000 rpm; leaf temperature, 28°C; relative humidity (RH), 68-70 %; [CO_2_], 100 μmol mol^−1^; PAR, 1500 μmol PAR m^−2^ s^−1^. Gas exchange measurements were automatically logged every minute and *g*_s_max was calculated as the average of the last 5 min at steady-state.

**Figure 1.**
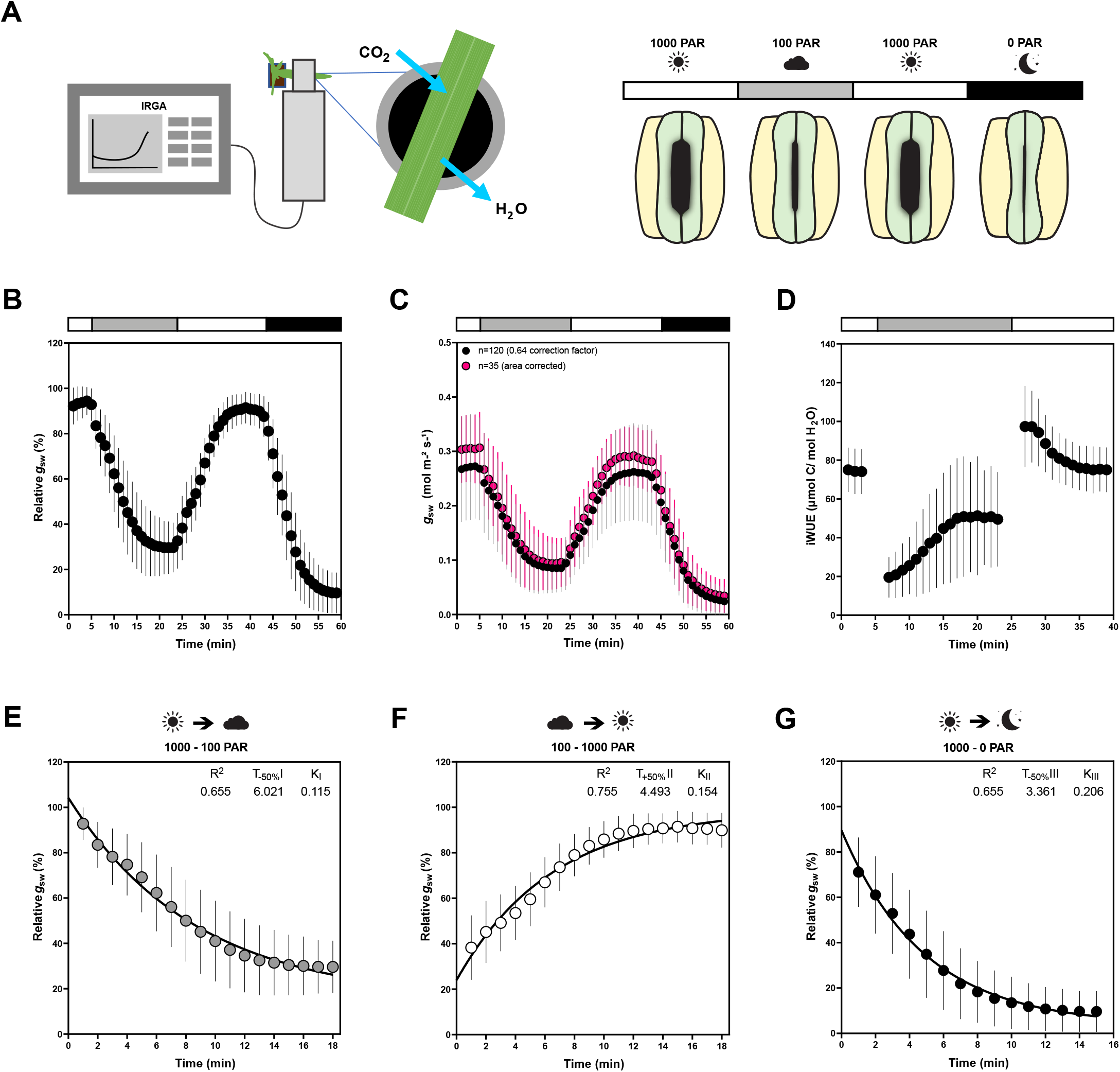
Leaf-level gas exchange measurements in response to changing light intensities reveal fast and consistent stomatal movements in *B. distachyon* Bd21-3. (A) Experimental setup for measuring leaf-level gas exchange parameters related with CO_2_ capture and H_2_O vapor loss by clamping a leaf in an IRGA chamber with controlled environmental conditions. Gas exchange is measured in changing light conditions (1000-100-1000-0 PAR) inducing stomatal closure in response to decreasing light intensity/darkness and stomatal opening in response to increasing light intensity. (B) Relative stomatal conductance (Rel *g*_sw_) during the light transitions (n=120, normalized to highest *g*_sw_ observed). (C) Absolute stomatal conductance (*g*_sw_) response to light transitions (in black, data from 120 corrected by average leaf area of 0.64 cm^2^ and in magenta data from a subset of 35 individuals corrected by individual leaf area). (D) Intrinsic water use efficiency (iWUE) response to light transitions (1000 - 100 - 100 PAR) (n=120, calculated as *A*/*g*_sw_). (E) One-phase decay exponential regression for the transition 1000 to 100 PAR (n = 120). (F) One-phase association exponential regression for the transition 100 to 1000 PAR (n = 120). (G) One-phase decay exponential regression for the transition 1000 to 0 PAR (n = 120). R^2^, half-time (T_±50%_), and rate constant (K) are indicated.

### Microscopy analysis of stomatal anatomical traits

Leaves of a subset of individuals (n= 4-6 per season; n=5 summer, n=6 in autumn, n=4 in winter) were collected after LI-6800 measurements and fixed in 7:1 ethanol:acetic acid. To prepare samples for imaging, leaf tissue was rinsed in water and mounted on slides in Hoyer’s solution. The abaxial side was imaged using a Leica DM5000B. For stomatal density (SD), 3-5 fields of view (0.290 mm^2^, 20X objective) per leaf were counted resulting in 60 - 160 stomata. For stomatal length (SL) and width (W_A_), stomata from 4-6 fields of view (0.0725 mm^2^, 40X objective) per leaf were measured resulting in 20 - 70 stomata per leaf.

### Correlation analysis and statistics

Correlation analysis for gas exchange parameters independent of leaf area like iWUE (ratio between *A* and *g*_sw_) and half-time of opening/closing were performed for all 120 individuals. Correlation analysis for gas exchange parameters dependent on leaf area like *g*_sw_ and *A* were only performed for the subset of 35 individuals, for which the accurate individual leaf area was determined. For the correlation analysis between steady-state gas exchange and environmental conditions, the last 5 min of steady-state gas exchange parameters (*A, g*_sw_ and iWUE) at the 2^nd^, 3^rd^ and 4^th^ light step (100, 1000 and 0 PAR) and their corresponding ambient conditions (𝒬_out_, RH and T) were averaged. Pearson correlation matrices represented as heatmaps were obtained for steady-state *g*_sw_, *A*, iWUE, 𝒬_out_, RH, T and time at 100, 1000 and 0 PAR. ROUT method was used to remove outliers (Motulsky & Brown, 2006).

For the correlation analysis between stomatal kinetics and steady-state *g*_sw_, pearson correlation matrices represented as heatmaps were obtained for steady-state initial/final *g*_sw_ and half-time (n=35). For the correlation between stomatal kinetics and environmental conditions, pearson correlation matrices represented as heatmaps were obtained for half-time, initial/final T, RH 𝒬_out_, and time of the day (n=120).

Relevant correlations between different pairs of parameters were represented with linear or non-linear regressions. Significant (p<0.05) and non-significant linear regressions were represented with solid and dashed lines, respectively. Non-linear regressions (quadratic function) were chosen for correlations with time of the day as we observe an axis of symmetry around noon, for which a non-linear model was biologically more appropriate.

For the correlation analysis between stomatal anatomical traits and growth environmental conditions, Pearson correlation matrices represented as heatmaps were obtained for SD, SL and environmental growth conditions (average T, average RH and day length). Pearson correlation matrices and linear regressions were obtained for the correlation analysis between stomatal anatomy and gas exchange parameters (steady-state *g*_sw_, *A*, iWUE, and T_50%_).

To test for significant differences between two groups we performed an unpaired t-test. One-way ANOVA followed by Tukey’s multiple comparison test was used when comparing more than 2 groups. P-values are indicated directly in the graphs. All analyses were performed on GraphPad Prism version 9.1.0, GraphPad Software, San Diego, California USA, www.graphpad.com.

### Stomatal morphometric analysis and anatomical gsmax calculations

To characterize fully open stomata, leaves were treated with 4 μM fusicoccin (Santa Cruz Biotechnology, Inc.) solution in opening-closing buffer (50 mM KCl, 10 mM MES-KOH). Collected leaves were dipped into 70 % ethanol and infiltrated with fusicoccin solution. For infiltration, a needleless syringe was used to infiltrate the leaf tissue on the adaxial side until the tissue was visibly wet. Infiltrated leaves were then cut into smaller pieces (approx. 3 - 5 mm long) and incubated overnight in fusicoccin solution in the light. To analyse closed stomata, leaves were treated with 50 μM ABA (Merck) solution in opening-closing buffer (50 mM KCl, 10 mM MES-KOH) as described for fusicoccin treatment and incubated overnight in the dark. Before imaging on the confocal microscope, leaves were stained with propidium iodide (1%). Z-stacks of 30 open and 30 closed stomata from 3 different individuals each were taken on the Leica TCS SP8 confocal microscope. The obtained z-stacks were analysed using Fiji. For each stoma, pore length (PL), pore width (PW) at the centre of the pore, guard cell length (GCL), right and left guard cell width at the middle of the stoma (GCW_C_) and stoma width at the apices were measured (W_A_) on the z sum projection image. To measure the exact pore area, each pore was manually traced with the polygon selection tool. To measure pore depth (l), the central pore part was selected with the rectangle selection tool and resliced starting at the top, avoiding interpolation. Pore depth was measured on the z sum projection of the reslice.

The leaves assessed for physiological *g*_s_max were fixed with ethanol:acetic acid 7:1. GCL, W_A_ and stomatal width at the centre (WC) were measured on light microscope pictures (20-40 stomata per leaf). GC width was calculated as half of W_A_ or W_C_. Stomatal density (SD) was obtained by counting the number of stomata in 5 different areas per leaf (20X objective).

For the anatomical maximum stomatal conductance calculations, we used the anatomical *g*_s_max equation from Franks & Farquhar (2001, see Fig. 5H), where SD is the stomatal density (stomata mm^−2,^), a_max_ is the maximum pore area (μm^2^), l is the pore depth (μm), d is the diffusivity of water in air (0.0000249 m^2^ s ^−1^, at 25°C), v is the molar volume of air (0.024464 m^3^ mol^−1^, at 25°C), and π is the mathematical constant. Maximum pore area was either measured by hand-tracing the stomatal pore of fully open stomata or calculated as an ellipse (with major axis equal to pore length and minor to half the pore length) or a rectangle (pore width x pore length).

## Results

### *B. distachyon* shows fast and consistent stomatal gas exchange in response to changing light

The plants’ physiology including stomatal gas exchange dynamics are strongly influenced by the environmental conditions the plant is exposed to (Arve et al., 2013; Durand et al., 2020; Matthews et al., 2017). Closed-system infra-red gas analyzers (IRGA) allow gas exchange measurements within a chamber with tightly controlled environmental settings regardless of ambient conditions (Douthe et al., 2018). However, plants have previously developed in and acclimated to a specific environment. Furthermore, during measurements, most distal parts of the plant remain exposed to ambient environmental conditions that might significantly differ from the conditions in the IRGA chamber (Fig. 1A). To quantify the consistency of stomatal responses and the influence of variable greenhouse conditions on gas exchange, we analyzed gas exchange parameters and kinetics of 120 wild-type *B. distachyon* individuals (Bd21-3) over the course of two years in a partially environmentally controlled greenhouse. The ambient conditions in the greenhouse varied remarkably over the course of the 120 IRGA measurements (Fig. S1A-C) and the measurements covered a broad range of hours of the day (time) from 6 am to 7 pm (Fig. S1D). We obtained consistent and reproducible stomatal light-responses (R^2^= 0.66-0.76) (Fig. 1B, E-G) despite variation in absolute stomatal conductance (*g*_sw_) levels (Fig. 1C).. Because the *B. distachyon* leaf is smaller than the 2 cm^2^ chamber used, *g*_sw_ was corrected using the average leaf area from a data subset for which we measured and corrected for the actual individual leaf area (n=35) (Fig. S1E), to obtain a mean approximation of gas exchange levels for the total 120 individuals. The data subset corrected with the actual leaf area (Fig. 1C, magenta dots) nicely overlapped with the average correction of the 120 individuals (Fig. 1C, black dots) and together revealed reasonable variation of absolute *g*_sw_ values. Importantly, the 35 leaf area corrected measurements covered the range of environmental conditions observed in all 120 individuals (Fig. S1A-D).

The first light transition (1000 to 100 PAR) resulted in a 70 % decrease in *g*_sw_ with a half-time of 6 minutes (T_-50%_I = 6.021 min) (Fig. 1E). An increase in light intensity (100 to 1000 PAR) induced an exponential increase in *g*_sw_ with a half-time of less than 5 min (T_+50%_II = 4.493 min) until reaching similar *g*_sw_ as in the previous high light step (1000 PAR) (Fig. 1F). Switching from 1000 to 0 PAR resulted in strikingly fast stomatal closure with a half-time of only ∼3 min (T_-50%_III = 3.361 min) and, thus, represented the quickest of the three light transition responses (Fig.1G). *g*_sw_ was on average 0.29 ± 0.06 mol m^−2^ s^−1^ at high light and 0.10 ± 0.05 mol m^−2^ s^−1^ at low light (Fig. 1C). We observed an average of 0.015 ± 0.013 mol m^−2^ s^−1^) of residual *g*_sw_ in darkness (Fig. 1C). At high light, iWUE was on average 77 ± 13 μmol CO_2_/mol H_2_O, whereas at low light iWUE was 51± 26 μmol CO_2_/mol H_2_O (Fig. 1 D). *A* was on average 21 ± 4 μmol m^−2^ s^−1^ at high light and 4 ± 1 μmol m^−2^ s^−1^ at low light (Fig. S1F).

Together, *B. distachyon* shows fast stomatal light responses typical for grasses, which were consistent over 120 measurements.

### Quantitative effects of greenhouse environmental fluctuations on gas exchange in *B. distachyon*

To quantify how the different environmental conditions affected gas exchange, we performed correlation analysis between gas exchange parameters (stomatal conductance (*g*_sw_), carbon assimilation (*A*) and intrinsic water-use efficiency (iWUE)) and environmental conditions (temperature (T), ambient light intensity (𝒬_out_), relative humidity (RH), time of the day (time)). Correlations were done separately for high light steady-state (1000 PAR, Fig 2A), low light steady-state (100 PAR, Fig. 2G), steady-state in darkness (0 PAR, Fig. S2A) and for opening and closing kinetics (Fig. S2I-K). Because exact leaf area was only measured for a subset of 35 individuals to calculate accurate absolute *g*_sw_ and *A*, the correlation analysis between environmental parameters and absolute gas exchange parameters (i.e. *g*_sw_ and *A*) was performed using the 35 individuals only. On the other hand, the 120 samples were used for correlation analysis between environmental conditions and leaf-area independent parameters like iWUE and stomatal kinetics (half-time).

**Figure 2.**
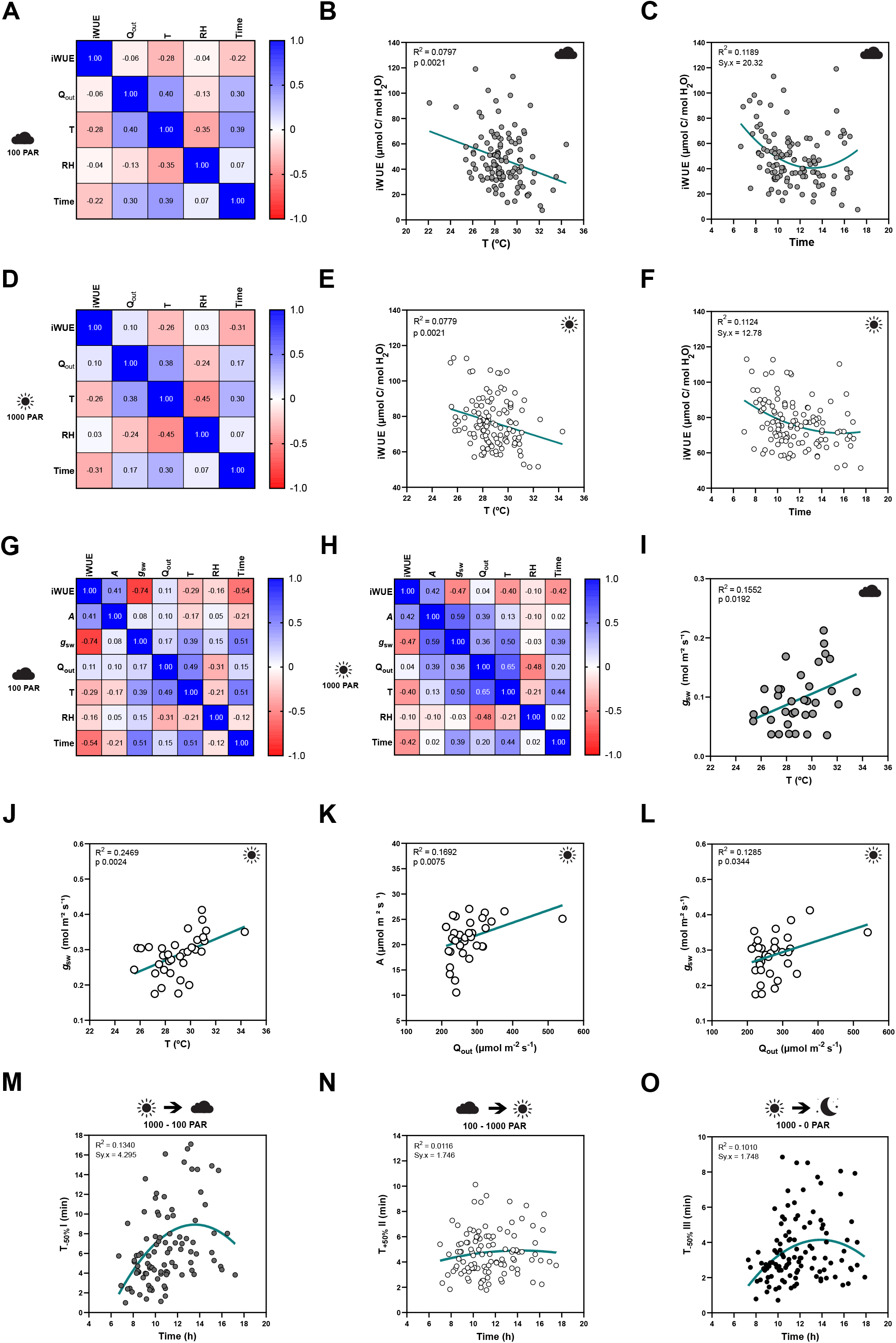
Influence of temperature, ambient light and circadian rhythm on leaf-level gas exchange. (A) Correlation matrix between iWUE and environment (Q_out_, T, RH and time) of 120 measurements of wild-type *B. distachyon* (Bd21-3) at 100 PAR (2^nd^ light step). (B) Linear regression between T and iWUE at 100 PAR (n=116). (C) Non-linear regression between time and iWUE at 100 PAR (n=116). (D) Correlation matrix between iWUE and environment (Q_out_, T, RH and time) of 120 measurements of wild-type *B. distachyon* (Bd21-3) at 1000 PAR (3^rd^ light step). (E) Linear regression between T and iWUE at 1000 PAR (n=119). (F) Non-linear regression between time and iWUE at 1000 PAR (n=120). (G) Cor-relation matrix between gas exchange parameters (*A, g*_sw_ and iWUE) and environment (Q_out_, T, RH and time) of the 35 measurements (corrected by individual leaf area) of wild-type *B. distachyon* (Bd21-3) at 100 PAR (2^nd^ light step). (H) Correlation matrix between gas exchange parameters (*A, g*_sw_ and iWUE) and environment (Q_out_, T, RH and time) of the 35 measurements (corrected by individual leaf area) of wild-type *B. distachyon* (Bd21-3) at 1000 PAR (3^rd^ light step). (I) Linear regression between T and *g*_sw_ at 100 PAR (n=35). (J) Linear regression between T and *g*_sw_ at 1000 PAR (n=35). (K) Linear regression between Q_out_ and *A* at 1000 PAR (n=35). (L) Linear regression between Q_out_ and *g*_sw_ at 1000 PAR (n=35). (M) Non-linear regression between half-time of the transition 1000-100 PAR (T_-50%_I) and time of the day (time) (n=99). (N) Non-linear regression between half-time of the transition 100-1000 PAR (T_+50%_II) and time of the day (time) (n =111). (O) Non-linear regression between half-time of the transition 1000 - 0 PAR (T_-50%_III) and time of the day (time) (n=111). R^2^ and Sy.x or p-values are indicated.

iWUE was negatively correlated with T at both light conditions 100 and 1000 PAR (Fig. 2A, D). Increasing temperatures significantly associated with decreasing iWUE values (Fig. 2B, E). iWUE also correlated with time in both light conditions (Fig. 2A, D). A quadratic relation can be observed between time and iWUE (Fig. 2C, F), particularly at low light, with the lowest iWUE reached at midday (Fig. 2C). Similar correlations between iWUE and T or time were observed in the data subset (n=35) (Fig. 2G, H).

Regarding the steady-state gas exchange parameters, T showed a considerable influence on *g*_sw_ at both low and high light (Fig. 2G, H) and *g*_sw_ significantly increased with rising ambient temperatures (Fig. 2I, J). 𝒬_out_, on the other hand, correlated with both *A* and *g*_sw_ at high light (Fig. 2H). Both *A* and *g*_sw_ significantly increased with increasing 𝒬_out_(Fig.2K, L). Together, this explained why iWUE is only correlated with T but not with 𝒬_out_.

Lastly, time significantly correlated with *g* at all light conditions and also with *A* at low light (Fig. 2G-H, Fig. S2A-E, G). No significant correlations were observed between ambient conditions (𝒬_out_, T or RH) and *g*_sw_ at 0 PAR (Fig. S2E-G), even though an influence of T on *g*_sw_ is suggested (Fig. S2G) as observed at 100 and 1000 PAR (Fig. 2I-J).

Finally, stomatal kinetics (i.e. half-time T_50%_) significantly depended on the initial and/or final steady-state *g*_sw_(Fig. S2H), which in turn were affected by the environment (see above). In addition, our data also suggested an influence of circadian rhythm (time) on stomatal closure speed (half-time T_50%_) as stomata seem to close slower at noon (Fig. 2M-O, Fig. S2I-K).

In summary, fluctuations in ambient conditions such as temperature and light intensity during measurements, and circadian rhythm influenced steady-state gas exchange parameters and/or stomatal kinetics within a strictly controlled IRGA leaf chamber, which highlights the relevance of considering systemic effects on stomatal physiology experiments.

### Seasonal changes in greenhouse growth conditions affect stomatal anatomical traits and gas exchange

While the artificial light intensity and light-darkness cycles are controlled, the contribution of ambient sunlight intensity and day length (DL), average temperature (T) and relative humidity (RH) vary among seasons in our greenhouse (Fig. S3A, D-F). Stomatal anatomical traits such as stomatal density (SD) and stomatal size are strongly influenced by environmental cues to which the plants are exposed to during development (Casson & Gray, 2008; Liu et al., 2018; Qi & Torii, 2018; Terfa et al., 2020). To test the plasticity and variability of stomatal anatomical traits in *B. distachyon* wild-type plants, we quantified SD and stomatal length (SL) as a proxy for stomatal size from 15 individuals grown in different seasons—summer (May-June, n=5), autumn (October-November, n=6) and winter (January-February, n=4)—and correlated these traits with growth conditions (T, RH and day length (Fig. 3A, S3A). In summer, SD was ∼40% higher than in winter (105.8±13.1 vs. 74.2±6.8 stomata per mm^2^) and SL reduced by ∼10% (24.6±0.7 vs. 26.8±0.3 μm) (Fig. S3A-C).

**Figure 3.**
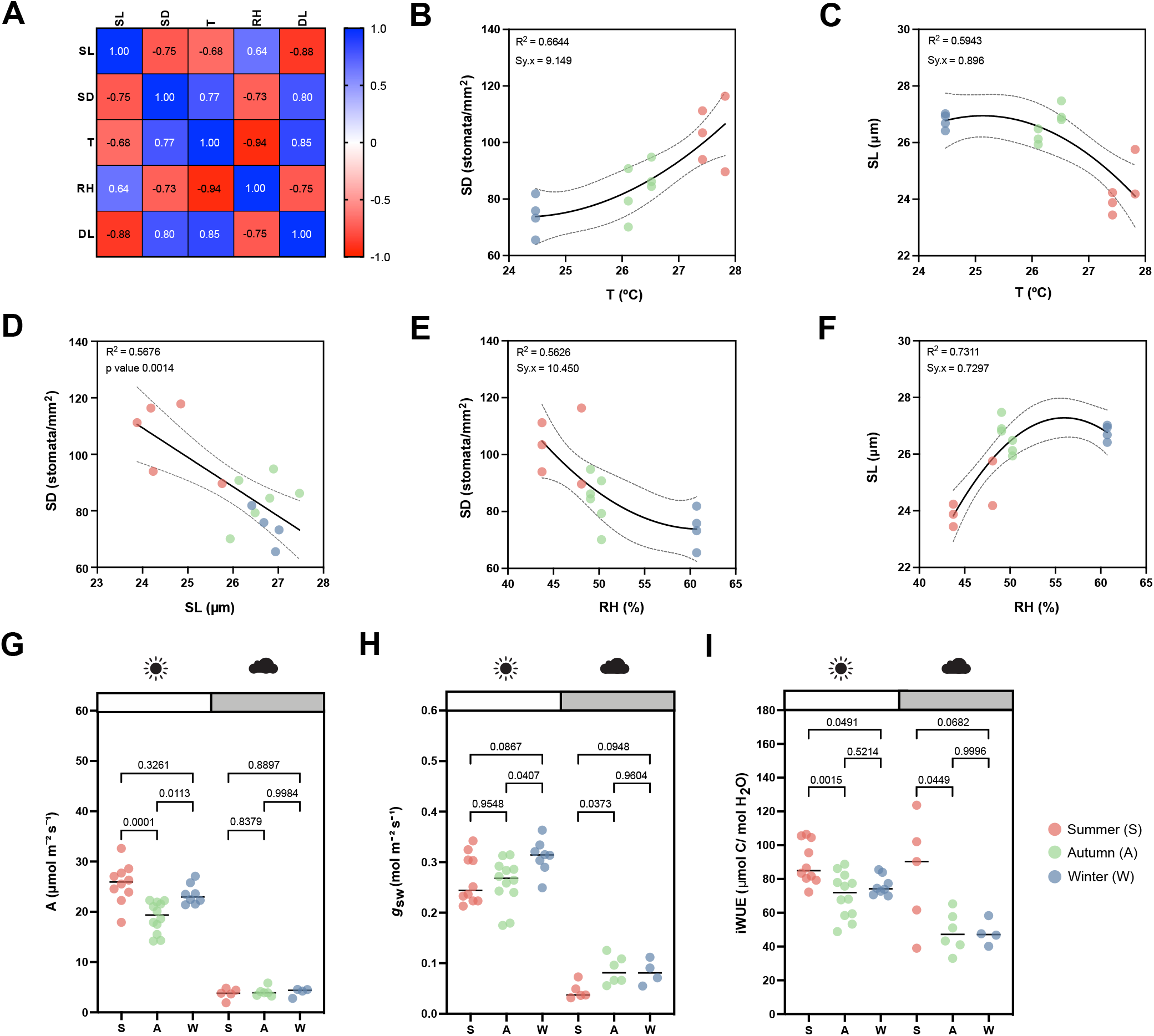
Effects of seasonal growth conditions on stomatal anatomical traits and on gas exchange. (A) Correlation matrix between stomatal length (SL), stomatal density (SD), average growth temperature (T), average growth relative humidity (RH) and day length (DL) (n=15). (B) Non-linear quadratic (second order polynomial regression) relation between T and SD (n=15). (C) Non-linear quadratic (second order polynomial regression) relation between T and SL (n=15). (D) Linear relation between SD and SL (n=15). (E) Non-linear quadratic (second order polynomial regression) relation between RH and SD (n = 15). (F) Non-linear quadratic (second order polynomial regression) relation between RH and SL (n=15). (G) Seasonal variation on *A* at 1000 (white) and 100 (grey) PAR (n=15). (H) Seasonal variation on *g*_sw_ at 1000 and 100 PAR (n=15). (I) Seasonal variation on iWUE at 1000 and 100 PAR (n=15). R^2^ and Sy.x or p-values are indicated.

Consequently and as previously described (Franks & Beerling, 2009; Haworth et al., 2021; Zhang et al., 2021), we observed a strong negative correlation between SD and SL (Fig. 3A, D) and a strong correlation between anatomy and environment (Fig. 3A). SD and SL correlated in an opposite manner with the different environmental parameters (Fig. 3A). An increase in T correlated with an increase in SD and a decrease in SL (Fig. 3B, C). Since T and RH were negatively correlated, RH correlated in an opposite manner with SD and SL (Fig. 3 E, F). SD and SL were also inversely correlated with DL, with longer days associated with shorter stomata and higher SD (Fig. S3G, H). Overall, summer plants grown during longer days with higher ambient light intensity, higher T, and lower RH (Fig. S3D-F), developed higher SD and lower SL (Fig. S3B, C). In autumn and winter, plants grown during shorter days with lower ambient light intensity, lower T and higher RH (Fig. S3D-F), developed lower SD and higher SL (Fig. S3B, C). However, which environmental parameter primarily caused changes to stomatal anatomy is unclear.

Besides a seasonal variation on stomatal anatomical traits, we observed seasonal variation on gas exchange (Fig. 3G-I). In summer, we observed higher *A* under high light than in autumn (Fig. 3G) but no significant difference in *g*_sw_(Fig. 3H). Consequently, iWUE was higher in summer than in autumn under high light conditions (Fig. 3I). We also observed an improvement in *A* from autumn to winter (Fig. 3G). Under low light conditions (100 PAR), no differences in *A* occurred among seasons (Fig. 3G). On the other hand, higher *g*_sw_ was observed in autumn and winter (Fig. 3H), resulting in lower iWUE at 100 PAR (Fig. 3I). When measuring physiological maximum stomatal conductance (*g*_s_max) of autumn/winter plants, the anatomical offset between SD and SL seemed to compensate for stomatal gas exchange maximum capacity, even though a non-significant decrease in average *g*_s_max was observed in autumn/winter (Fig. S3I). In conclusion, stomatal anatomical traits of wild-type *B. distachyon* are surprisingly plastic and variable among seasons even in a semi-controlled growth environment, likely contributing to the seasonal variation on functional traits.

### Stomatal anatomical traits influence gas exchange

Due to the seasonal variation in stomatal anatomy and functional traits, we quantified how the anatomical variation translates into changes in functional traits such as steady-state gas exchange parameters (at high light 1000 PAR, low light 100 PAR and darkness 0 PAR) and stomatal kinetics (Fig. S4).

In terms of steady-state *g*_sw_, lower SD (and higher SL, to a lesser extent) are the anatomical traits associated with higher operational stomatal conductance (*g*_sw_) in *B. distachyon* (Fig. 4A, B). Similarly, an increase in SD and a decrease in SL resulted in an increase of *A* at high light, while no effect was observed in 100 PAR (light limiting condition) (Fig.4C-D). Consequently, higher SD and lower SL result in higher iWUE (Fig. 4E, F), even though SD had a stronger effect on iWUE (Fig. 4F). Thus, the higher iWUE observed in summer (Fig. 3I) might be primarily caused by the higher SD and lower SL observed in this season (Fig. S3B, C). The correlations between anatomical traits (SD and SL) and *g*_sw_ were stronger in low light than in high light (Fig. S4, Fig. 4A, B) likely contributing to the higher seasonal variation in iWUE at low light than in high light (Fig. 3I).

**Figure 4.**
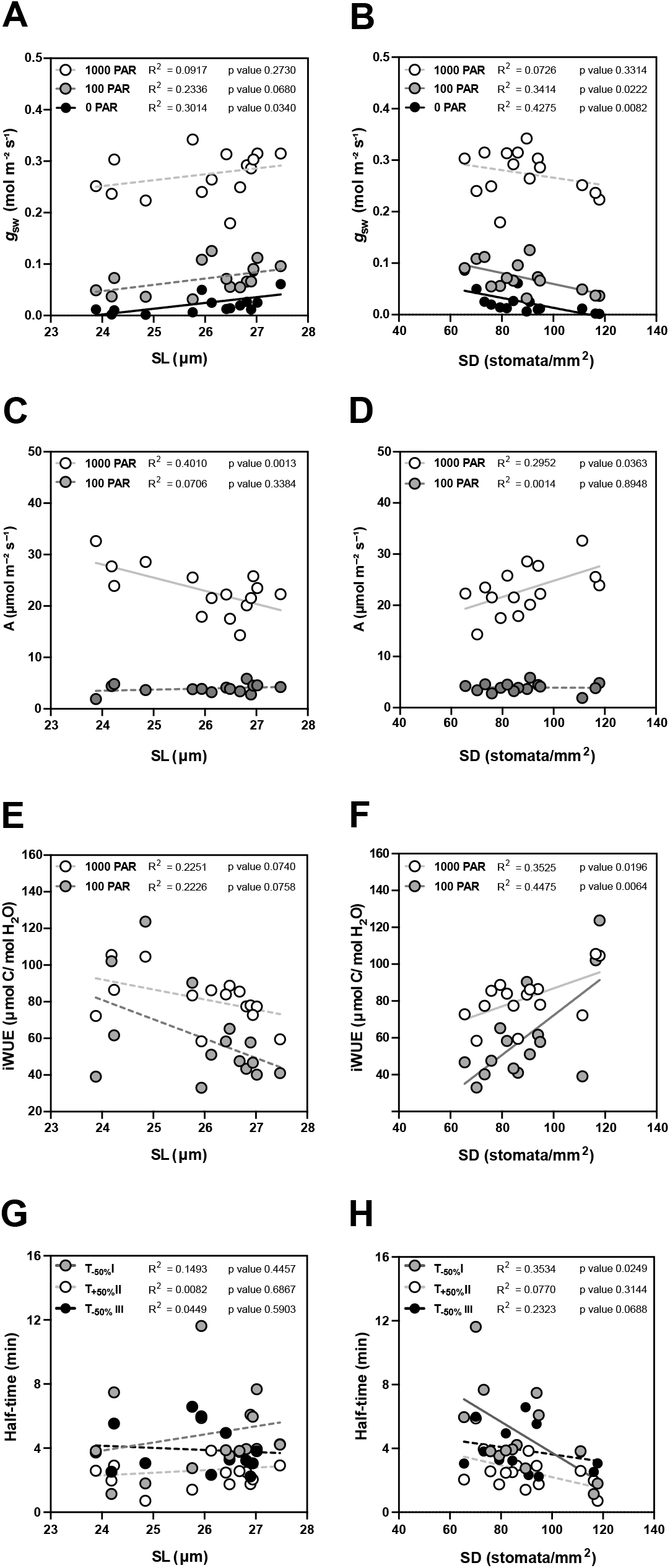
Impact of stomatal anatomical traits on steady-state gas exchange and stomatal kinetics. (A) Linear regressions between stomatal length (SL) and *g*_sw_ at 1000 (white dots), 100 (grey dots) and 0 (black dots) PAR (n =15). (B) Linear regressions between stomatal density (SD) and *g*_sw_ at 1000 (white dots), 100 (grey dots) and 0 (black dots) PAR (n =15). (C) Linear regressions between SL and *A* at 1000 (white dots) and 100 PAR (grey dots). (D) Linear regression between SD and *A* at 1000 (white) and 100 PAR (grey dots). (E) Linear regressions between SL and iWUE at 1000 PAR (white dots) and 100 PAR (grey dots) (n=15). (F) Linear regressions between SD and iWUE at 1000 PAR (white dots) and 100 PAR (grey dots) (n=15). (G) Linear regressions between SL and half-time (T_50%_) of the light transitions 1000-100 (grey dots), 100-1000 (white dots) and 1000-0 (black dots) PAR (n=15). (H) Linear regressions between SD and T_50%_ of the light transitions 1000-100 (grey dots), 100-1000 (white dots) and 1000-0 (black dots) PAR (n=15). R^2^ and p-values are indicated. Dashed lines indicate statistically non-significant linear regressions (p>0.05).

Regarding stomatal kinetics (T_50%_), the effect of SL on stomatal closure and opening was non-significant (Fig. 4G, Fig. S4). However, while the influence of SD on stomatal opening was also weak (Fig. 4H, Fig. S4), surprisingly stronger correlations and significant effects were observed between SD and stomatal closure kinetics (T_-50%_) (Fig. 4H, Fig.S4). High SD was strongly correlated with water-use efficiency as the increase of SD led to higher steady-state iWUE (Fig. 4F) and faster stomatal closure (Fig. 4H) contributing to higher water-use efficiency in changing environments. Overall, stomatal anatomy is strongly correlated with stomatal functioning and the seasonal variation in stomatal anatomy strongly contributed to seasonal variation in gas exchange.

### Morphometric analysis of graminoid stomata to optimize anatomical *g*_s_max predictions in *B. distachyon*

Finally, we wanted to mathematically describe the impact of the observed trade-off between SD and SL on maximum stomatal conductance (*g*_s_max) by calculating the theoretical anatomical *g*_s_max, which is based on the anatomical traits stomatal density (SD), maximum pore area (a_max_) and pore depth (l). While SD is assessed for any species simply by counting stomata per leaf area, formulae to calculate maximum pore area (a_max_, μm^2^) and pore depth (l, μm) were optimized for *Arabidopsis*-like stomatal morphologies and ellipsoid pores (Dow et al., 2014; Franks & Farquhar, 2001) (Fig. 5A).

**Figure 5.**
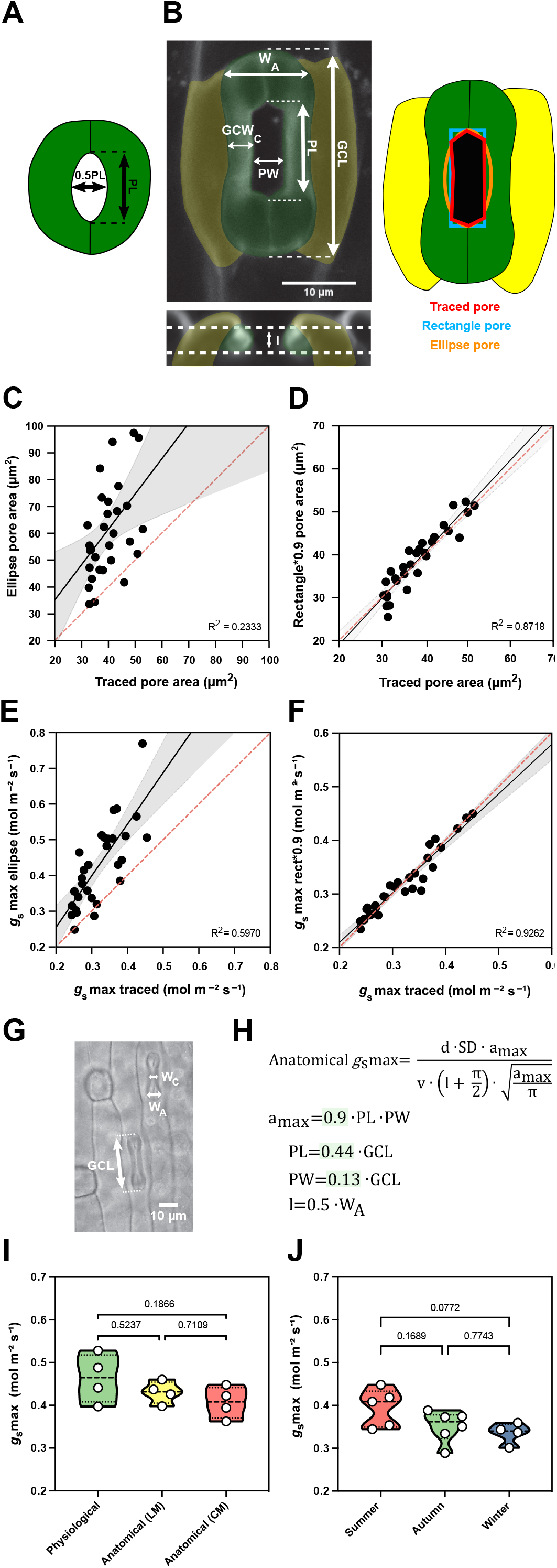
Morphometric analysis of graminoid *B. distachyon* stomata significantly improves anatomical *g*_s_max predictions. (A) *Arabidopsis*-like stoma and ellipse pore shape. (B) *B. distachyon* stomatal morphology traits measured; guard cell length (GCL), pore length (PL), pore width (PW), guard-cell width at the center of the stomata (GCW_C_), stomatal width at the apices (W_A_) and pore depth (l). Pore area hand-traced (red) or geometrically defined as an ellipse (orange) or a rectangle (blue). (C) Linear relation between hand-traced pore area and ellipse pore area. (D) Linear relation between hand-traced pore area and rectangle multiplied by 0.9. (E) Linear relation of anatomical *g*_s_max calculated with hand-traced pore area and with ellipse pore. (F) Linear relation of anatomical *g*_s_max calculated with hand-traced pore area and with rectangle pore multiplied by 0.9. (G) Anatomical parameters measured using light microscopy; stomatal width at the apex (W_A_) and guard cell length (GCL). (H) Anatomical maximum stomatal conductance (anatomical *g*_s_max) equation as defined by Franks & Farquhar (2001) and *B. distachyon* adjustments to calculate a_max_(0.9*PL*PW), PL (0.44*GCL), PW (0.13*GCL) and l (W_A_*0.5). (I) Comparison between physiological *g*_s_max, anatomical *g*_s_max (light microscopy, LM) and anatomical *g*_s_max (confocal microscopy, CM). (J) Comparison of anatomical *g*_s_max calculated for summer, autumn and winter plants with stomatal anatomical traits represented in Fig. S3B,C. R^2^ and p-values are indicated.

High-resolution confocal stacks of fusicoccin (fus)-treated open stomatal complexes in *B. distachyon* revealed hexagonal rather than elliptical pores (Fig. 5B). We performed careful morphometric analysis of open and closed stomata to characterize guard cell length (GCL), pore length (PL), pore width (PW), GC width at the centre (GCW_C_) and stomatal width at the apices (W_A_) (Fig. 5B). Furthermore, we manually traced and measured pore areas of fus-treated complexes to empirically determine a more appropriate way to calculate a_max_ of graminoid stomata. Calculating a_max_ as for *Arabidopsis-* like stomata (ellipse with major axis equal to pore length and minor to half the pore length, Fig. 5A) caused significant pore area overestimation compared to the manually traced pores of fus-treated open complexes (Fig. 5B, C). A rectangular rather than an ellipsoid approach still overestimated pore area (Fig. 5B, Fig. S5A), as the manually measured pore areas approximated 90±7% of the rectangular pore area (Fig. S5C). Thus, *B. distachyon* stomatal pore area can be accurately estimated by using rectangular pore area calculations multiplied by a correction factor of 0.9 (Fig. 5D).

Regarding pore depth (l), we observed that the mean pore depth measured from orthogonal resliced confocal stacks (3.37±0.4μm) approximates the mean GC width at the center (3.23±0.3μm) but not the GC width at the apices (5.92±0.4μm) in fully opened stomata (fus-treated) (Fig. S5G). In closed stomata (ABA-treated), on the other hand, the mean GC width at the centre was 2.48±0.3μm while the GC width at the apices was 3.52±0.6μm (Fig. S5G). Therefore, if not exactly measured from orthogonal sections, then pore depth could be approximated as central GC width of fully open stomata or apical GC width of closed stomata.

We then calculated anatomical *g*_s_max using hand-traced a_max_ and estimated a_max_ using formulae for i) ellipse, ii) rectangle pore and iii) rectangle pore multiplied by the correction factor 0.9. Pore depth was measured from orthogonal resliced confocal stacks (Fig. 5B) and stomatal density was determined by counting stomata in 3-5 different fields of view using light microscopy. We could observe that anatomical *g*_s_max using hand-traced a_max_ nicely correlated with anatomical *g*_s_max calculated for the rectangle pore multiplied by the correction factor 0.9 (Fig. 5F). This was not the case when using ellipse a_max_(Fig. 5E) or rectangular pore without the correction factor (Fig. S5B, C).

To determine a_max_ from simple light microscopy pictures, where pores are hard to see, we calculated correction coefficients to estimate pore length and width from GC length. By calculating the ratios of the morphometrically determined GCL, PL, and PW, we found that PL is 44±3% of GCL and PW is 13±3% of GCL (Fig. S5D-F). Thus, for calculations using light microscopy pictures, we estimated PL as 0.44*GCL and PW as 0.13*GCL (Fig. 5G, H). To approximate pore depth (l), GC width at the apices was used for closed or partially open stomata (½ of the stoma width at the apices) (Fig. 5G, H, Fig. S5G)) and GC width at the centre was used for fully open stomata.

Next, we tested if our adjustments for anatomical *g*_s_max calculations could be used to reliably predict physiological *g*_s_max in *B. distachyon*. We performed IRGA-based measurements of *g*_s_max (physiological *g*_s_max) in four independent individuals, collected these exact leaf zones, and measured anatomical traits from segments of those by using both standard light microscopy (after fixation with ethanol:acetic acid 7:1) and confocal microscopy (after treatment with fusicoccin). No significant differences were found between physiological *g*_s_max and anatomical *g*_s_max based on measured anatomical parameters using light microscopy (LM) or confocal microscopy (CM) (Fig. 5I, Fig. S5H). In summary, the optimized formula for accurate anatomical *g*_s_max estimation can be used to reliably predict physiological *g*_s_max in *B. distachyon*.

Finally, we calculated anatomical *g*_s_max for the summer, autumn and winter individuals whose anatomical traits (SD and SL) were shown in Figure S3B, C. Even though a decrease in average anatomical *g*_s_max was observed in autumn and winter, this difference was non-significant (Fig. 5J). These results match our observations in physiological *g*_s_max measurements between summer and autumn/winter plants (Fig. S3I). Ultimately, these findings suggest that the trade-off between SD and SL in wild-type *B. distachyon* might serve as a mechanism to maximize stomatal conductance capacity in different environments.

## Discussion

Consistent and reproducible stomatal kinetics were observed for *B. distachyon* regardless of the variable greenhouse conditions. *B. distachyon* displayed the fast stomatal movements typical for grass species, which are faster than most non-grass species with kidney-shaped GC (Franks & Farquhar, 2007; Grantz & Assmann, 1991; McAusland et al., 2016; Merilo et al., 2014). The quick stomatal movements of grasses like *B. distachyon* are associated with the graminoid morphology, where two lateral subsidiary cells (SCs) flank the central, dumbbell-shaped GCs (Gray et al., 2020; Nunes et al., 2020; Stebbins & Shah, 1960). Fast stomatal movements in grasses require SCs (Raissig et al., 2017), which might function as specialized ion reservoirs (Raschke & Fellows, 1971) and mechanically accommodate GC movement to accelerate both stomatal opening and closing (Franks & Farquhar, 2007). In addition, the reduced volume-to-surface ratio of dumbbell-shaped GCs likely requires less exchange of water and ions to be pressurized (Franks & Farquhar, 2007).

Furthermore, no major asymmetry between closure and opening speed was observed. This is consistent with the results from a comparison of stomatal kinetics between eight species with kidney-shaped GCs and seven species with dumbbell-shaped GCs, where species with dumbbell-shaped GCs displayed the quickest responses and showed the most similarity between opening and closure times (McAusland et al., 2016). Nonetheless, the fastest stomatal movement in *B. distachyon* was dark-induced stomatal closure (1000 to 0 PAR). Faster stomatal closing than opening has been previously described for several species and suggested to be a water-conserving strategy (Lawson & Vialet-Chabrand, 2019; Leakey et al., 2019; McAusland et al., 2016). Fast stomatal movements are important to quickly adjust stomatal pores to avoid excess of water vapor loss through stomata (*g*_sw_) during suboptimal carbon assimilation (*A*) in low light. Intrinsic water use efficiency (iWUE, the ratio between *A* and *g*_sw_), varied between 50 and 80 μmol mol^−1^ which is consistent with the range of iWUE described for other C_3_ grass species such as wheat (25-65 μmol mol^−1^) and rice (50-80 μmol mol^−1^) (Giuliani et al., 2013; Jahan et al., 2014).

Despite the consistent stomatal responsiveness in *B. distachyon*, we observed a significant influence of the time of the day on light-response stomatal kinetics. While diurnal variation on gas exchange has been well described for C_3_ species (Matthews et al., 2017; Miao et al., 2021; Resco de Dios, 2017; Roussel et al., 2009; Stangl et al., 2019; Vahisalu et al., 2008), the observed diurnal variation on stomatal responsiveness to changing light in the tightly regulated conditions of the IRGA chambers was compelling. Stomatal closure and opening speed were mainly affected by the time of the day and by steady-state *g*_sw_ prior and/or after the change in light intensity. A similar dependence of *g*_sw_ kinetics on light intensity transitions, on the time of the day and on steady-state *g*_sw_ prior to light intensity changes has been recently described in *Musa spp*. (Eyland et al., 2021).

Steady-state *g*_sw_, on the other hand, was significantly influenced by the environmental conditions. Stomatal conductance was stimulated by increasing ambient temperatures. This effect has been observed and described as a leaf cooling mechanism to cope with higher temperatures (Gommers, 2020; Lamba et al., 2018; Sonawane et al., 2021; Urban et al., 2017b, 2017a). Yet, after exceeding a certain threshold, such high temperatures may lead to stress-induced stomatal closure (Faria et al., 1996; Ikkonen et al., 2015; Yamori et al., 2006; Zhou et al., 2015).. In addition, increasing ambient light intensity also significantly impacted IRGA measurements by triggering increases in *g*_sw_ and *A* levels. Recent studies have also reported systemic stomatal responses to light and heat in Arabidopsis (Devireddy et al., 2018, 2020) and in response to darkness and elevated CO_2_ in birch and poplar (Ehonen et al., 2020). Consistently, our results suggest that *B. distachyon* stomata integrate both local environmental cues and systemic signals from distal parts of the plant. Therefore, it is important to monitor environmental conditions and consider their impact on gas exchange measurements, particularly in greenhouse or field studies with significant environmental fluctuations.

Apart from the effect of environmental conditions on gas exchange, different growth conditions had a major impact on anatomical traits. Stomatal density (SD) and length (SL) inversely varied among seasons due to significant variation in ambient growth conditions. The seasonal trade-off between SD and SL mostly maintained maximum gas exchange capacity in the different growth environments. In addition, higher SD was associated with faster stomatal closure and the combination of higher SD and lower SL associated with improved iWUE in wild-type *B. distachyon*. This suggests that higher SD and lower SL, which are associated with improved stomatal responsiveness and more water use efficient gas exchange, are a morphological adaptation to summer. Higher stomatal densities in warmer environments were reported and associated as an ecophysiological significant response for leaf evaporative cooling (Carlson et al., 2016; Hill et al., 2015). In contrast, autumn and winter seasons feature shorter days, decreased light intensity and colder temperatures, which can negatively affect photosynthesis (Feng et al., 2018; Yamasaki et al., 2002). Thus, a decrease in stomatal density to increase the leaf surface allocated to light harvesting, compensated by an increase in stomatal size to maintain maximum gas exchange capacity, might sustain optimal gas exchange and photosynthesis in winter. While we observed higher iWUE in *B. distachyon* wild-type plants with lower SL and higher SD, crop species (wheat, barley and rice) overexpressing an inhibitor of stomatal development (*EPF1*) show a reduction in SD and improved iWUE (Caine et al., 2018; Dunn et al., 2019; Hughes et al., 2017). Thus, genetically modifying SD (and/or SL) in *B. distachyon* beyond the intraspecific range of variation might allow to improve iWUE. *B. distachyon* genotypes varying in single morphological traits may help to better understand the independent influence of SL and SD on gas exchange kinetics, capacity and water-use efficiency, and verify the correlations observed in this present study.

Some studies described a negative correlation between stomatal size and stomatal speed (e.g. (Drake et al., 2013; Durand et al., 2020; Kardiman & Ræbild, 2018). In a study comparing different rice genotypes, stomatal size was negatively correlated with stomatal half-time (Zhang et al., 2019), with larger stomata being faster. In a comparison of 15 different species (including 7 grass species), however, smaller stomata were associated with faster stomatal movements (McAusland et al., 2016). Other approaches using a broader range of species from different plant groups (by comparing 7 (Elliott-Kingston et al., 2016) and 31 (Haworth et al., 2018) different species) suggest that stomatal speed is not related to SL but rather positively correlated with SD, as we observed in *B. distachyon*. A major effect of SD and minor effect of SL on stomatal speed under fluctuating light has also been described in a study comparing different *Arabidopsis* genotypes (Sakoda et al., 2020). Higher SD could potentially cause proximity effects triggering stomata in close vicinity to react to local changes in a more coordinated manner. However, the effect of the variation of SL and SD on gas exchange speediness may vary among species and/or genotypes. Nonetheless, environmentally-induced SD and SL variation and its impact on gas exchange must be considered during long term studies performed in greenhouse or field settings.

To facilitate correlations of anatomical stomatal traits to theoretical gas exchange maximum capacity (anatomical *g*_s_max) in *B. distachyon*, we adjusted established equations (Dow et al., 2014; Franks & Farquhar, 2001) to the graminoid stomatal morphology. For grass stomata, stomatal pores are rather hexagons than ellipses and the equation presented in this study accurately predicted anatomical *g*_s_max for *B. distachyon*. Therefore, it can be reliably used to predict gas exchange capacity of *B. distachyon* genotypes varying in anatomical traits. Differences between the anatomical and physiological *g*_s_max might reveal impaired stomatal signalling and thus, provide a tool to identify mutant phenotypes of stomatal function. In addition, anatomical *g*_s_max could help to weigh the effect of variations in single morphological traits (e.g. stomatal density, stomatal size, pore area) on *g*_s_max.

In conclusion, stomatal conductance kinetics are fast and consistent in the grass model species *B. distachyon*. Nevertheless and even though stomata primarily respond to the local environment (i.e. within the IRGA chamber), ambient light intensity, temperature and time of the day can have systemic effects on gas exchange influencing results from IRGA measurements. Stomatal anatomical traits are highly plastic and environmental-responsive and, furthermore, have a major impact on gas exchange. For that reason, the effect of growth conditions on stomatal anatomical traits must be considered in leaf-level gas exchange studies.

## Acknowledgments

The authors would like to thank Michael Schilbach for managing the greenhouse and gardening support. Furthermore, the authors acknowledge all members of the lab for carefully reading and critically evaluating this manuscript. This work was supported by a DAAD Study Scholarship (to MWS) and by the German Research Foundation (DFG) Emmy Noether Programme Grant RA 3117/1-1 (to MTR).

## Data and Coding Availability

All gas exchange data and anatomical data used in this study can be found in Supplementary Dataset 1. Stomatal images are available upon request.

## References

Arve, L. E., Terfa, M. T., Gislerød, H. R., Olsen, J. E., & Torre, S. (2013). High relative air humidity and continuous light reduce stomata functionality by affecting the ABA regulation in rose leaves. Plant, Cell Environment, 36(2), 382–392. https://doi.org/10.1111/j.1365-3040.2012.02580.x

Bertolino, L. T., Caine, R. S., & Gray, J. E. (2019). Impact of Stomatal Density and Morphology on Water-Use Efficiency in a Changing World. Frontiers in Plant Science, 10, 225. https://doi.org/10.3389/fpls.2019.00225

Caine, R. S., Yin, X., Sloan, J., Harrison, E. L., Mohammed, U., Fulton, T., Biswal, A. K., Dionora, J., Chater, C. C., Coe, R. A., Bandyopadhyay, A., Murchie, E. H., Swarup, R., Quick, W. P., & Gray, J. E. (2018). Rice with reduced stomatal density conserves water and has improved drought tolerance under future climate conditions. The New Phytologist. https://doi.org/10.1111/nph.15344

Carlson, J. E., Adams, C. A., & Holsinger, K. E. (2016). Intraspecific variation in stomatal traits, leaf traits and physiology reflects adaptation along aridity gradients in a South African shrub. Annals of Botany, 117(1), 195–207. https://doi.org/10.1093/aob/mcv146

Casson, S., & Gray, J. E. (2008). Influence of environmental factors on stomatal development. The New Phytologist, 178(1), 9–23. https://doi.org/10.1111/j.1469-8137.2007.02351.x

Devireddy, A. R., Arbogast, J., & Mittler, R. (2020). Coordinated and rapid whole-plant systemic stomatal responses. The New Phytologist, 225(1), 21–25. https://doi.org/10.1111/nph.16143

Devireddy, A. R., Zandalinas, S. I., Gómez-Cadenas, A., Blumwald, E., & Mittler, R. (2018). Coordinating the overall stomatal response of plants: Rapid leaf-to-leaf communication during light stress. Science Signaling, 11(518). https://doi.org/10.1126/scisignal.aam9514

Douthe, C., Gago, J., Ribas-Carbó, M., Núñez, R., Pedrol, N., & Flexas, J. (2018). Measuring Photosynthesis and Respiration with Infrared Gas Analysers. In A.M. Sánchez-Moreiras & M. J. Reigosa (Eds.), Advances in Plant Ecophysiology Techniques (pp. 51–75). Springer International Publishing. https://doi.org/10.1007/978-3-319-93233-0_4

Dow, G. J., Berry, J. A., & Bergmann, D. C. (2014). The physiological importance of developmental mechanisms that enforce proper stomatal spacing in Arabidopsis thaliana. The New Phytologist, 201(4), 1205–1217. https://doi.org/10.1111/nph.12586

Drake, P. L., Froend, R. H., & Franks, P. J. (2013). Smaller, faster stomata: scaling of stomatal size, rate of response, and stomatal conductance. Journal of Experimental Botany, 64(2), 495–505. https://doi.org/10.1093/jxb/ers347

Dunn, J., Hunt, L., Afsharinafar, M., Al Meselmani, M., Mitchell, A., Howells, R., Wallington, E., Fleming, A. J., & Gray, J. E. (2019). Reduced stomatal density in bread wheat leads to increased water-use efficiency. Journal of Experimental Botany. https://doi.org/10.1093/jxb/erz248

Durand, M., Brendel, O., Buré, C., & Le Thiec, D. (2020). Changes in irradiance and vapour pressure deficit under drought induce distinct stomatal dynamics between glasshouse and field-grown poplars. The New Phytologist, 227(2), 392–406. https://doi.org/10.1111/nph.16525

Ehonen, S., Holtta, T., & Kangasjärvi, J. (2020). Systemic signaling in the regulation of stomatal conductance. Plant Physiology. https://doi.org/10.1104/pp.19.01543

Elliott-Kingston, C., Haworth, M., Yearsley, J. M., Batke, S. P., Lawson, T., & McElwain, J. C. (2016). Does Size Matter? Atmospheric CO2 May Be a Stronger Driver of Stomatal Closing Rate Than Stomatal Size in Taxa That Diversified under Low CO2. Frontiers in Plant Science, 7, 1253. https://doi.org/10.3389/fpls.2016.01253

Engineer, C. B., Hashimoto-Sugimoto, M., Negi, J., Israelsson-Nordström, M., Azoulay-Shemer, T., Rappel, W.-J., Iba, K., & Schroeder, J. I. (2016). CO2 Sensing and CO2 Regulation of Stomatal Conductance: Advances and Open Questions. Trends in Plant Science, 21(1), 16–30. https://doi.org/10.1016/j.tplants.2015.08.014

Eyland, D., van Wesemael, J., Lawson, T., & Carpentier, S. (2021). The impact of slow stomatal kinetics on photosynthesis and water use efficiency under fluctuating light. Plant Physiology. https://doi.org/10.1093/plphys/kiab114

Faralli, M., Matthews, J., & Lawson, T. (2019). Exploiting natural variation and genetic manipulation of stomatal conductance for crop improvement. Current Opinion in Plant Biology, 49, 1–7. https://doi.org/10.1016/j.pbi.2019.01.003

Faria, T., García-Plazaola, J. I., Abadía, A., Cerasoli, S., Pereira, J. S., & Chaves, M. M. (1996). Diurnal changes in photoprotective mechanisms in leaves of cork oak (Quercus suber) during summer. Tree Physiology, 16(1_2), 115–123. https://doi.org/10.1093/treephys/16.1-2.115

Feng, L., Raza, M. A., Li, Z., Chen, Y., Khalid, M. H. B., Du, J., Liu, W., Wu, X., Song, C., Yu, L., Zhang, Z., Yuan, S., Yang, W., & Yang, F. (2018). The Influence of Light Intensity and Leaf Movement on Photosynthesis Characteristics and Carbon Balance of Soybean. Frontiers in Plant Science, 9, 1952. https://doi.org/10.3389/fpls.2018.01952

Franks, P. J., & Beerling, D. J. (2009). Maximum leaf conductance driven by CO2 effects on stomatal size and density over geologic time. Proceedings of the National Academy of Sciences of the United States of America, 106(25), 10343–10347. https://doi.org/10.1073/pnas.0904209106

Franks, P. J., Drake, P. L., & Beerling, D. J. (2009). Plasticity in maximum stomatal conductance constrained by negative correlation between stomatal size and density: an analysis using Eucalyptus globulus. Plant, Cell & Environment, 32(12), 1737–1748. https://doi.org/10.1111/j.1365-3040.2009.02031.x

Franks, P. J., & Farquhar, G. D. (2001). The effect of exogenous abscisic acid on stomatal development, stomatal mechanics, and leaf gas exchange in Tradescantia virginiana. Plant Physiology, 125(2), 935–942. https://doi.org/10.1104/pp.125.2.935

Franks, P. J., & Farquhar, G. D. (2007). The mechanical diversity of stomata and its significance in gas-exchange control. Plant Physiology, 143(1), 78–87. https://doi.org/10.1104/pp.106.089367

Giuliani, R., Koteyeva, N., Voznesenskaya, E., Evans, M. A., Cousins, A. B., & Edwards, G. E. (2013). Coordination of Leaf Photosynthesis, Transpiration, and Structural Traits in Rice and Wild Relatives (Genus Oryza). Plant Physiology, 162(3), 1632–1651. https://doi.org/10.1104/pp.113.217497

Gommers, C. (2020). Keep Cool and Open Up: Temperature-Induced Stomatal Opening [Review of Keep Cool and Open Up: Temperature-Induced Stomatal Opening]. Plant Physiology, 182(3), 1188–1189. https://doi.org/10.1104/pp.20.00158

Grantz, D. A., & Assmann, S. M. (1991). Stomatal response to blue light: water use efficiency in sugarcane and soybean*. Plant, Cell & Environment, 14(7), 683–690. https://doi.org/10.1111/j.1365-3040.1991.tb01541.x

Gray, A., Liu, L., & Facette, M. (2020). Flanking Support: How Subsidiary Cells Contribute to Stomatal Form and Function. Frontiers in Plant Science, 11, 881. https://doi.org/10.3389/fpls.2020.00881

Haworth, M., Marino, G., Loreto, F., & Centritto, M. (2021). Integrating stomatal physiology and morphology: evolution of stomatal control and development of future crops. In Oecologia. https://doi.org/10.1007/s00442-021-04857-3

Haworth, M., Scutt, C. P., Douthe, C., Marino, G., Gomes, M. T. G., Loreto, F., Flexas, J., & Centritto, M. (2018). Allocation of the epidermis to stomata relates to stomatal physiological control: Stomatal factors involved in the evolutionary diversification of the angiosperms and development of amphistomaty. Environmental and Experimental Botany, 151, 55–63. https://doi.org/10.1016/j.envexpbot.2018.04.010

Hill, K. E., Guerin, G. R., Hill, R. S., & Watling, J. R. (2015). Temperature influences stomatal density and maximum potential water loss through stomata of Dodonaea viscosa subsp. angustissima along a latitude gradient in southern Australia. Australian Journal of Botany, 62(8), 657–665. https://www.publish.csiro.au/bt/bt14204

Hughes, J., Hepworth, C., Dutton, C., Dunn, J. A., Hunt, L., Stephens, J., Waugh, R., Cameron, D. D., & Gray, J. E. (2017). Reducing Stomatal Density in Barley Improves Drought Tolerance without Impacting on Yield. Plant Physiology, 174(2), 776–787. http://www.plantphysiol.org/lookup/doi/10.1104/pp.16.01844

Ikkonen, E. N., Shibaeva, T. G., Rosenqvist, E., & Ottosen, C. O. (2015). Daily temperature drop prevents inhibition of photosynthesis in tomato plants under continuous light. Photosynthetica, 53(3), 389–394. https://doi.org/10.1007/s11099-015-0115-4

Jahan, E., Amthor, J. S., Farquhar, G. D., Trethowan, R., & Barbour, M. M. (2014). Variation in mesophyll conductance among Australian wheat genotypes. Functional Plant Biology: FPB, 41(6), 568–580. https://doi.org/10.1071/FP13254

Jezek, M., & Blatt, M. R. (2017). The Membrane Transport System of the Guard Cell and Its Integration for Stomatal Dynamics. Plant Physiology, 174(2), 487–519. https://doi.org/10.1104/pp.16.01949

Kardiman, R., & Ræbild, A. (2018). Relationship between stomatal density, size and speed of opening in Sumatran rainforest species. Tree Physiology, 38(5), 696–705. https://doi.org/10.1093/treephys/tpx149

Kollist, H., Nuhkat, M., & Roelfsema, M. R. G. (2014). Closing gaps: linking elements that control stomatal movement. The New Phytologist, 203(1), 44–62. https://doi.org/10.1111/nph.12832

Lamba, S., Hall, M., Räntfors, M., Chaudhary, N., Linder, S., Way, D., Uddling, J., & Wallin, G. (2018). Physiological acclimation dampens initial effects of elevated temperature and atmospheric CO2 concentration in mature boreal Norway spruce. Plant, Cell & Environment, 41(2), 300–313. https://doi.org/10.1111/pce.13079

Lawson, T., & Blatt, M. R. (2014). Stomatal size, speed, and responsiveness impact on photosynthesis and water use efficiency. Plant Physiology, 164(4), 1556–1570. https://doi.org/10.1104/pp.114.237107

Lawson, T., & Matthews, J. (2020). Guard Cell Metabolism and Stomatal Function. Annual Review of Plant Biology. https://doi.org/10.1146/annurev-arplant-050718-100251

Lawson, T., & Vialet-Chabrand, S. (2019). Speedy stomata, photosynthesis and plant water use efficiency. The New Phytologist, 221(1), 93–98. https://doi.org/10.1111/nph.15330

Leakey, A. D. B., Ferguson, J. N., Pignon, C. P., Wu, A., Jin, Z., Hammer, G. L., & Lobell, D. B. (2019). Water Use Efficiency as a Constraint and Target for Improving the Resilience and Productivity of C3 and C4 Crops. Annual Review of Plant Biology, 70, 781–808. https://doi.org/10.1146/annurev-arplant-042817-040305

Liu, C., He, N., Zhang, J., Li, Y., Wang, Q., Sack, L., & Yu, G. (2018). Variation of stomatal traits from cold temperate to tropical forests and association with water use efficiency. Functional Ecology, 32(1), 20–28. https://doi.org/10.1111/1365-2435.12973

Matthews, J. S. A., Vialet-Chabrand, S. R. M., & Lawson, T. (2017). Diurnal Variation in Gas Exchange: The Balance between Carbon Fixation and Water Loss. Plant Physiology, 174(2), 614–623. https://doi.org/10.1104/pp.17.00152

McAusland, L., Vialet-Chabrand, S., Davey, P., Baker, N. R., Brendel, O., & Lawson, T. (2016). Effects of kinetics of light-induced stomatal responses on photosynthesis and water-use efficiency. The New Phytologist, 211(4), 1209–1220. https://doi.org/10.1111/nph.14000

Merilo, E., Jõesaar, I., Brosché, M., & Kollist, H. (2014). To open or to close: species-specific stomatal responses to simultaneously applied opposing environmental factors. The New Phytologist, 202(2), 499–508. https://doi.org/10.1111/nph.12667

Miao, Y., Cai, Y., Wu, H., & Wang, D. (2021). Diurnal and Seasonal Variations in the Photosynthetic Characteristics and the Gas Exchange Simulations of Two Rice Cultivars Grown at Ambient and Elevated CO2. In Frontiers in Plant Science (Vol. 12). https://doi.org/10.3389/fpls.2021.651606

Motulsky, H. J., & Brown, R. E. (2006). Detecting outliers when fitting data with nonlinear regression - a new method based on robust nonlinear regression and the false discovery rate. BMC Bioinformatics, 7, 123. https://doi.org/10.1186/1471-2105-7-123

Murata, Y., Mori, I. C., & Munemasa, S. (2015). Diverse stomatal signaling and the signal integration mechanism. Annual Review of Plant Biology, 66(1), 369–392. http://www.annualreviews.org/doi/10.1146/annurev-arplant-043014-114707

Nunes, T. D. G., Zhang, D., & Raissig, M. T. (2020). Form, development and function of grass stomata. The Plant Journal: For Cell and Molecular Biology, 101(4), 780–799. https://doi.org/10.1111/tpj.14552

Qi, X., & Torii, K. U. (2018). Hormonal and environmental signals guiding stomatal development. BMC Biology, 16(1), 21. https://doi.org/10.1186/s12915-018-0488-5

Raissig, M. T., Matos, J. L., Anleu Gil, M. X., Kornfeld, A., Bettadapur, A., Abrash, E., Allison, H. R., Badgley, G., Vogel, J. P., Berry, J. A., & Bergmann, D. C. (2017). Mobile MUTE specifies subsidiary cells to build physiologically improved grass stomata. Science, 355(6330), 1215–1218. https://doi.org/10.1126/science.aal3254

Raschke, K., & Fellows, M. P. (1971). Stomatal movement in Zea mays: Shuttle of potassium and chloride between guard cells and subsidiary cells. Planta, 101(4), 296–316. https://doi.org/10.1007/BF00398116

Resco de Dios, V. (2017). Circadian Regulation and Diurnal Variation in Gas Exchange. Plant Physiology, 175(1), 3–4. https://doi.org/10.1104/pp.17.00984

Roussel, M., Dreyer, E., Montpied, P., Le-Provost, G., Guehl, J.-M., & Brendel, O. (2009). The diversity of 13C isotope discrimination in a Quercus robur full-sib family is associated with differences in intrinsic water use efficiency, transpiration efficiency, and stomatal conductance. In Journal of Experimental Botany (Vol. 60, Issue 8, pp. 2419–2431). https://doi.org/10.1093/jxb/erp100

Sakoda, K., Yamori, W., Shimada, T., Sugano, S. S., Hara-Nishimura, I., & Tanaka, Y. (2020). Higher Stomatal Density Improves Photosynthetic Induction and Biomass Production in Arabidopsis Under Fluctuating Light. Frontiers in Plant Science, 11, 589603. https://doi.org/10.3389/fpls.2020.589603

Sierla, M., Waszczak, C., Vahisalu, T., & Kangasjärvi, J. (2016). Reactive Oxygen Species in the Regulation of Stomatal Movements. Plant Physiology, 171(3), 1569–1580. https://doi.org/10.1104/pp.16.00328

Sonawane, B. V., Koteyeva, N. K., Johnson, D. M., & Cousins, A. B. (2021). Differences in leaf anatomy determines temperature response of leaf hydraulic and mesophyll CO 2 conductance in phylogenetically related C 4 and C 3 grass species. In New Phytologist. https://doi.org/10.1111/nph.17287

Stangl, Z. R., Tarvainen, L., Wallin, G., Ubierna, N., Räntfors, M., & Marshall, J. D. (2019). Diurnal variation in mesophyll conductance and its influence on modelled water-use efficiency in a mature boreal Pinus sylvestris stand. Photosynthesis Research, 141(1), 53–63. https://doi.org/10.1007/s11120-019-00645-6

Stebbins, G. L., & Shah, S. S. (1960). Developmental Studies of Cell Differentiation in the Epidermis of Monocotyledons. II. Cytological Features of Stomatal Development in the Gramineae. Developmental Biology, 2, 477–500.

Terfa, M. T., Olsen, J. E., & Torre, S. (2020). Blue Light Improves Stomatal Function and Dark-Induced Closure of Rose Leaves (Rosa x hybrida) Developed at High Air Humidity [Data set]. https://doi.org/10.3389/fpls.2020.01036

Urban, J., Ingwers, M., McGuire, M. A., & Teskey, R. O. (2017a). Stomatal conductance increases with rising temperature. Plant Signaling & Behavior, 12(8), e1356534. https://doi.org/10.1080/15592324.2017.1356534

Urban, J., Ingwers, M. W., McGuire, M. A., & Teskey, R. O. (2017b). Increase in leaf temperature opens stomata and decouples net photosynthesis from stomatal conductance in Pinus taeda and Populus deltoides x nigra. Journal of Experimental Botany, 68(7), 1757–1767. https://doi.org/10.1093/jxb/erx052

Vahisalu, T., Kollist, H., Wang, Y.-F., Nishimura, N., Chan, W.-Y., Valerio, G., Lamminmäki, A., Brosché, M., Moldau, H., Desikan, R., Schroeder, J. I., & Kangasjärvi, J. (2008). SLAC1 is required for plant guard cell S-type anion channel function in stomatal signalling. In Nature (Vol. 452, Issue 7186, pp. 487–491). https://doi.org/10.1038/nature06608

Yamasaki, T., Yamakawa, T., Yamane, Y., Koike, H., Satoh, K., & Katoh, S. (2002). Temperature acclimation of photosynthesis and related changes in photosystem II electron transport in winter wheat. Plant Physiology, 128(3), 1087–1097. https://doi.org/10.1104/pp.010919

Yamori, W., Suzuki, K., Noguchi, K. O., Nakai, M., & Terashima, I. (2006). Effects of Rubisco kinetics and Rubisco activation state on the temperature dependence of the photosynthetic rate in spinach leaves from contrasting growth temperatures. In Plant, Cell and Environment (Vol. 29, Issue 8, pp. 1659–1670). https://doi.org/10.1111/j.1365-3040.2006.01550.x

Zhang, L., Wang, S., Yang, X., Cui, X., & Niu, H. (2021). An Intrinsic Geometric Constraint on Morphological Stomatal Traits. Frontiers in Plant Science, 12, 628. https://doi.org/10.3389/fpls.2021.658702

Zhang, Q., Peng, S., & Li, Y. (2019). Increase rate of light-induced stomatal conductance is related to stomatal size in the genus Oryza. Journal of Experimental Botany, 70(19), 5259–5269. https://doi.org/10.1093/jxb/erz267

Zhou, H., Xu, M., Pan, H., & Yu, X. (2015). Leaf-age effects on temperature responses of photosynthesis and respiration of an alpine oak, Quercus aquifolioides, in southwestern China. Tree Physiology, 35(11), 1236–1248. https://doi.org/10.1093/treephys/tpv101

