## Supplemental Figures and Dataset for "Quantitative effects of environmental variation on stomatal anatomy and gas exchange in a grass model"

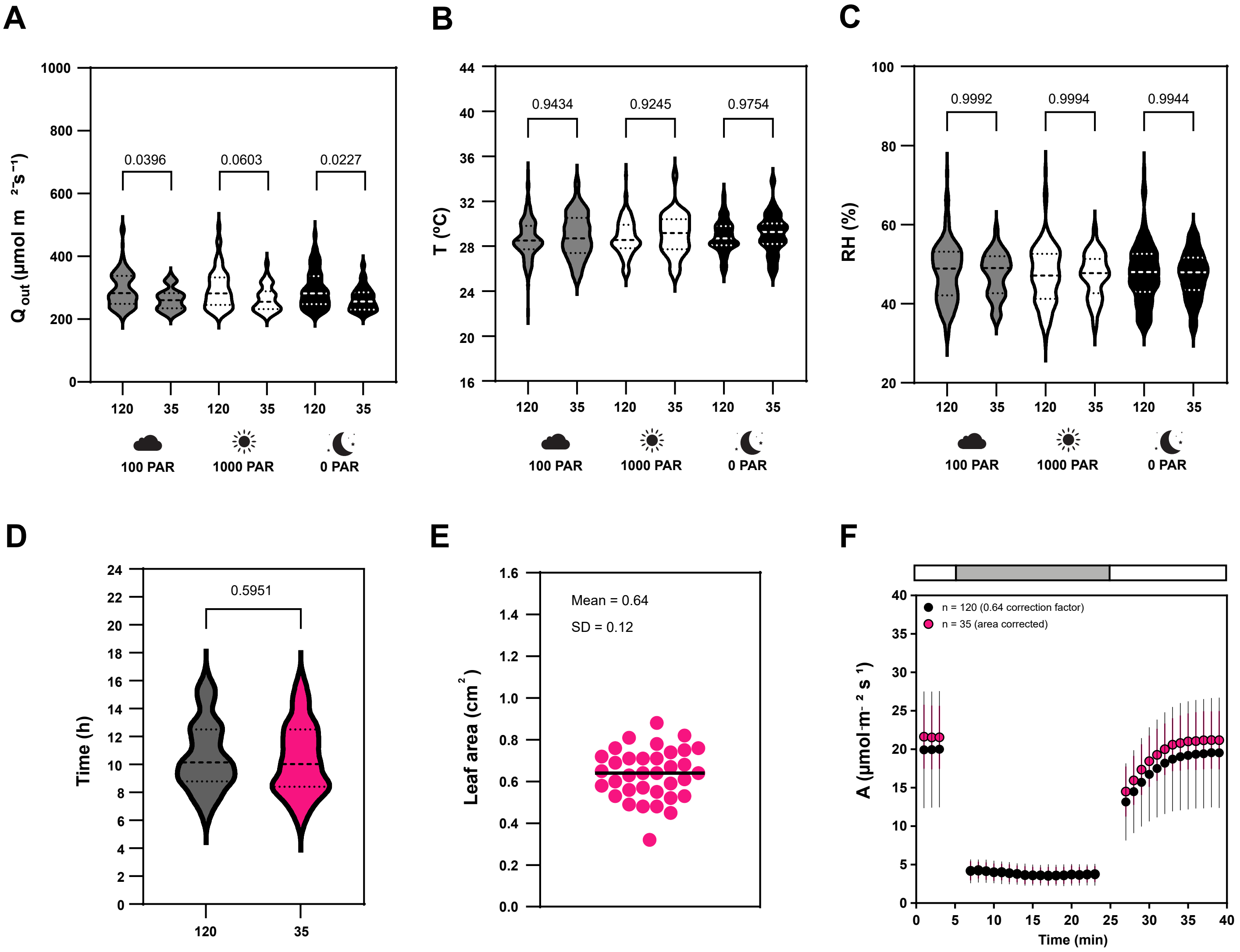


**Figure S1 - Environmental conditions in the greenhouse, leaf area and carbon assimilation.** (A) Ambient light during LI-6800 measurements (last 5 min of each steady-state used for correlation analysis at 100, 1000 and 0 PAR; both the total set of 120 measurements and the area-corrected subset of 35 individuals are shown). (B) Temperature during LI-6800 measurements (last 5 min of each steady-state used for correlation analysis 100, 1000 and 0 PAR; both the total set of 120 measurements and the area-corrected subset of 35 individuals are shown). (C) Relative humidity during LI-6800 measurements (last 5 min of each steady-state used for correlation analysis 100, 1000 and 0 PAR; both the total set of 120 measurements and the area-corrected subset of 35 individuals are shown). (D) Range of starting time of Li-6800 measurements of the 120 individuals (grey) and of the data subset of 35 individuals (magenta). (E) Leaf area accurately measured in a subset of 35 individuals (0.64±0.12). (F) Carbon assimilation during the light transitions 1000-100-1000 PAR (in black, data from 120 corrected by average leaf area of 0.64 $cm^2$ and in magenta data from a subset of 35 individuals corrected by individual leaf area). P-values from t-tests are indicated.

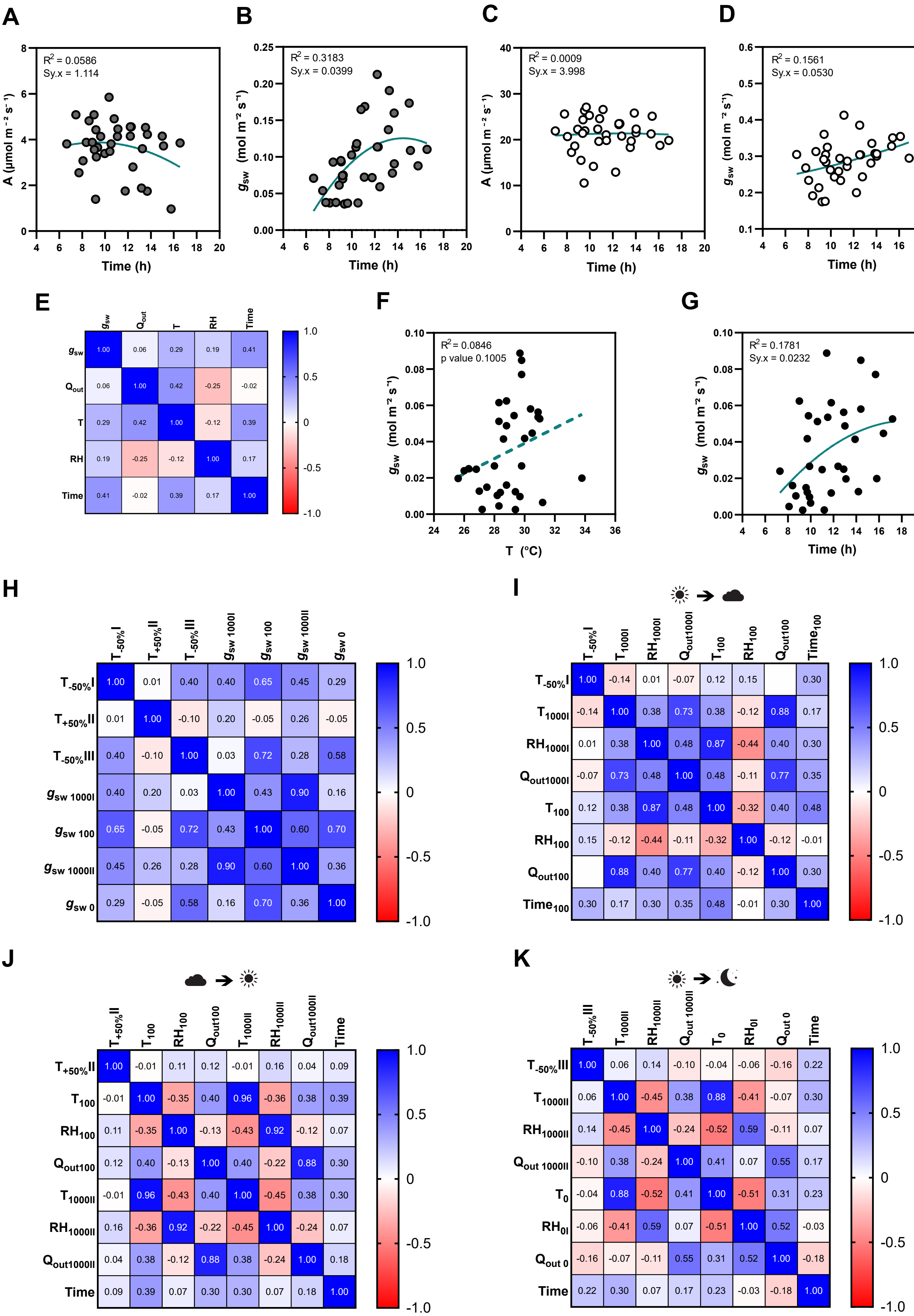
A
R2 = 0.0586
Sy.x = 1.114
A (µmol m⁻² s⁻¹)
Time (h)
B
R2 = 0.3183
Sy.x = 0.0399
gsw (mol m⁻² s⁻¹)
Time (h)
C
R2 = 0.0009
Sy.x = 3.998
A (µmol m⁻² s⁻¹)
Time (h)
D
R2 = 0.1561
Sy.x = 0.0530
gsw (mol m⁻² s⁻¹)
Time (h)
E
gsw Qout T RH Time
gsw 1.00 0.06 0.29 0.19 0.41
Qout 0.06 1.00 0.42 -0.25 -0.02
T 0.29 0.42 1.00 -0.12 0.39
RH 0.19 -0.25 -0.12 1.00 0.17
Time 0.41 -0.02 0.39 0.17 1.00
F
R2 = 0.0846
p value 0.1005
gsw (mol m⁻² s⁻¹)
T (°C)
G
R2 = 0.1781
Sy.x = 0.0232
gsw (mol m⁻² s⁻¹)
Time (h)
H
T-50%I T+50%II T-50%III gsw 1000I gsw 100 gsw 1000II gsw 0
T-50%I 1.00 0.01 0.40 0.40 0.65 0.45 0.29
T+50%II 0.01 1.00 -0.10 0.20 -0.05 0.26 -0.05
T-50%III 0.40 -0.10 1.00 0.03 0.72 0.28 0.58
gsw 1000I 0.40 0.20 0.03 1.00 0.43 0.90 0.16
gsw 100 0.65 -0.05 0.72 0.43 1.00 0.60 0.70
gsw 1000II 0.45 0.26 0.28 0.90 0.60 1.00 0.36
gsw 0 0.29 -0.05 0.58 0.16 0.70 0.36 1.00
I
T-50%I T1000I RH1000I Qout1000I T100 RH100 Qout100 Time100
T-50%I 1.00 -0.14 0.01 -0.07 0.12 0.15 0.30
T1000I -0.14 1.00 0.38 0.73 0.38 -0.12 0.88 0.17
RH1000I 0.01 0.38 1.00 0.48 0.87 -0.44 0.40 0.30
Qout1000I -0.07 0.73 0.48 1.00 0.48 -0.11 0.77 0.35
T100 0.12 0.38 0.87 0.48 1.00 -0.32 0.40 0.48
RH100 0.15 -0.12 -0.44 -0.11 -0.32 1.00 -0.12 -0.01
Qout100 0.88 0.40 0.77 0.40 -0.12 1.00 0.30
Time100 0.30 0.17 0.30 0.35 0.48 -0.01 0.30 1.00
J
T+50%II T100 RH100 Qout100 T1000II RH1000II Qout1000II Time
T+50%II 1.00 -0.01 0.11 0.12 -0.01 0.16 0.04 0.09
T100 -0.01 1.00 -0.35 0.40 0.96 -0.36 0.38 0.39
RH100 0.11 -0.35 1.00 -0.13 -0.43 0.92 -0.12 0.07
Qout100 0.12 0.40 -0.13 1.00 0.40 -0.22 0.88 0.30
T1000II -0.01 0.96 -0.43 0.40 1.00 -0.45 0.38 0.30
RH1000II 0.16 -0.36 0.92 -0.22 -0.45 1.00 -0.24 0.07
Qout1000II 0.04 0.38 -0.12 0.88 0.38 -0.24 1.00 0.18
Time 0.09 0.39 0.07 0.30 0.30 0.07 0.18 1.00
K
T-50%III T1000II RH1000II Qout 1000II T0 RH0I Qout 0 Time
T-50%III 1.00 0.06 0.14 -0.10 -0.04 -0.06 -0.16 0.22
T1000II 0.06 1.00 -0.45 0.38 0.88 -0.41 -0.07 0.30
RH1000II 0.14 -0.45 1.00 -0.24 -0.52 0.59 -0.11 0.07
Qout 1000II -0.10 0.38 -0.24 1.00 0.41 0.07 0.55 0.17
T0 -0.04 0.88 -0.52 0.41 1.00 -0.51 0.31 0.23
RH0I -0.06 -0.41 0.59 0.07 -0.51 1.00 0.52 -0.03
Qout 0 -0.16 -0.07 -0.11 0.55 0.31 0.52 1.00 -0.18
Time 0.22 0.30 0.07 0.17 0.23 -0.03 -0.18 1.00

**Figure S2 - Effect of environmental conditions on gas exchange and stomatal kinetics.** (A) Non-linear relation between *A* and time at 100 PAR (n=35). (B) Non-linear relation between $g_{sw}$ and time at 100 PAR (n=35). (C) Non-linear relation between *A* and time at 1000 PAR (n=35). (D) Non-linear relation between $g_{sw}$ and time at 1000 PAR (n=35). (E) Correlation matrix between $g_{sw}$ at 0 PAR and environmental conditions ($Q_{out}$, T, RH and time) (n=35). (F) Linear relation between $g_{sw}$ and $Q_{out}$ at 0 PAR (n=33). (G) Non-linear relation between $g_{sw}$ and time at 0 PAR (n=33). (H) Correlation matrix between stomatal half-time of the light transitions 1000-100 PAR($T_{-50\%}$I), 100-1000 PAR ($T_{+50\%}$II) and 1000-0 PAR ($T_{-50\%}$III) and initial/final step steady-state $g_{sw}$ (n=35). (I) Correlation matrix of stomatal closure half-time during the first light transition from 1000 to 100 PAR ($T_{-50\%}$I), T (1000 and 100 PAR), RH (1000 and 100 PAR), $Q_{out}$(1000 and 100 PAR) and time (n=120). (J) Correlation matrix of stomatal opening half-time during the second light transition from 100 to 1000 PAR ($T_{+50\%}$II), T (100 and 1000 PAR), RH (100 and 1000 PAR), $Q_{out}$(100 and 1000 PAR) and time (n=120). (K) Correlation matrix of stomatal closure half-time during the third light transition from 1000 to 0 PAR ($T_{+50\%}$II), T (1000 and 0 PAR), RH (1000 and 0 PAR), $Q_{out}$(1000 and 0 PAR), time (n=120). $R^2$, and Sy.x or p-values are indicated.

**A**

| | Summer | | | | | Autumn | | | | | Winter | | | | |
|---|---|---|---|---|---|---|---|---|---|---|---|---|---|---|---|
| | SL | SD | T | RH | DL | SL | SD | T | RH | DL | SL | SD | T | RH | DL |
| Mean | 24,58 | 105,8 | 27,58 | 45,47 | 16,00 | 26,62 | 84,28 | 26,32 | 49,67 | 10,00 | 26,76 | 74,15 | 24,47 | 60,72 | 9,000 |
| SD | 0,7462 | 13,10 | 0,2191 | 2,372 | 0,000 | 0,5579 | 8,750 | 0,2246 | 0,6627 | 0,000 | 0,2762 | 6,799 | 0,000 | 0,000 | 0,000 |

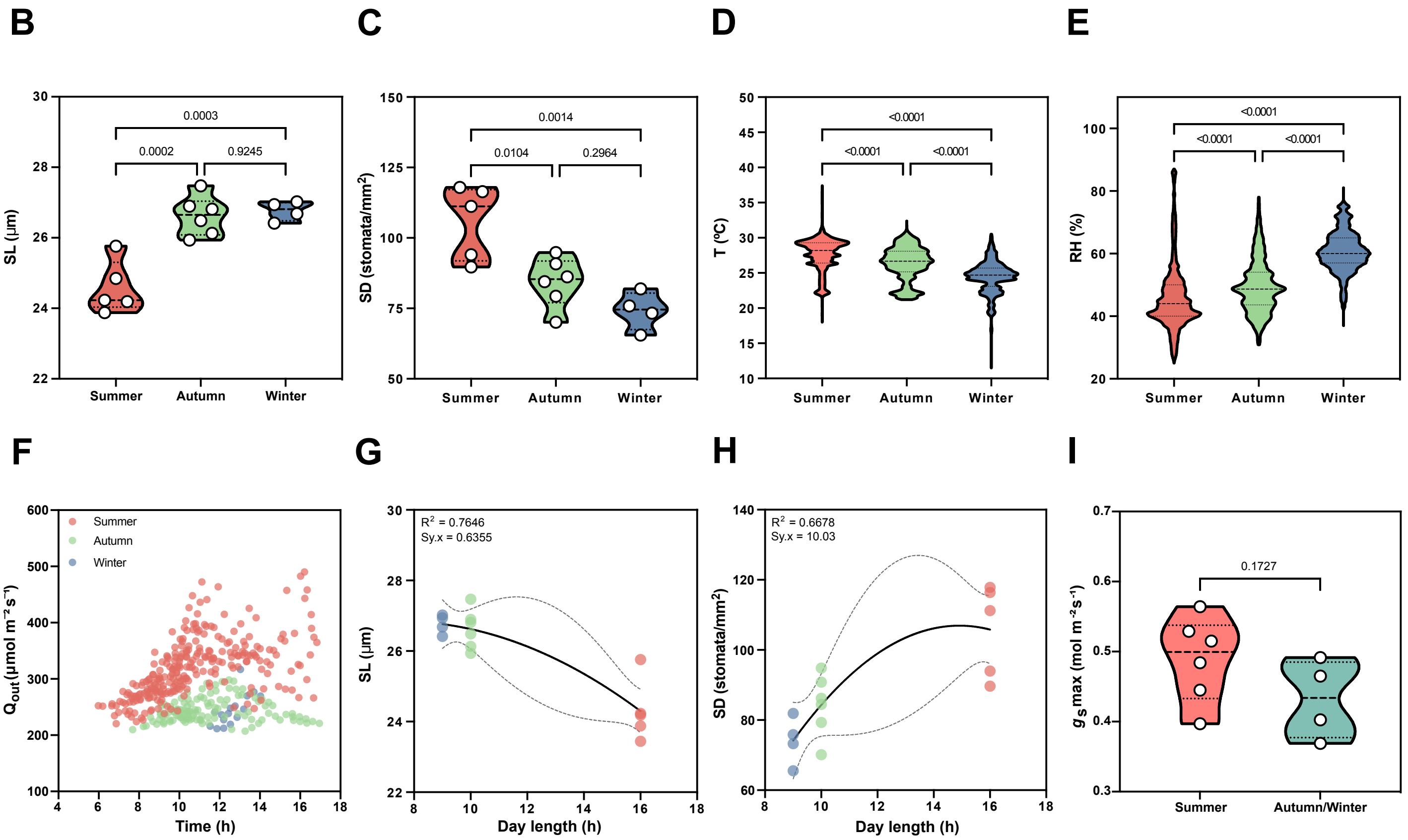


**Figure S3 - Seasonal effects on stomatal anatomy and environmental conditions.** (A) Mean values and standard deviation of stomatal length (SL), stomatal density (SD), average growth temperature (T), average growth relative humidity (RH) and day length (DL) for the summer, autumn and winter plants. (B) SL seasonal variation (summer n=5, autumn n=6, winter n=4). (C) SD seasonal variation (summer n=5, autumn n=6, winter n=4).). (D) Growth temperature variation for each season 2 weeks before leaf collection. (E) Growth relative humidity variation for each season 2 weeks before leaf collection. (F) Daily variation in ambient light intensity (data from LI-6800 measurements). (G) Non-linear relation between SL and day length. (H) Non-linear relation between SD and day length. (I) Physiological maximum stomatal conductance ($g_s$max) of summer (n=6) and autumn/winter (n=3/n=1, respectively) plants. The $g_s$max measurements were not performed on the plants that were anatomically assessed, which is why the number of individuals varies (summer n=6, fall n=3, winter n=1) and why autumn and winter measurements were combined. $R^2$ and Sy.x or p-values are indicated.

**A**

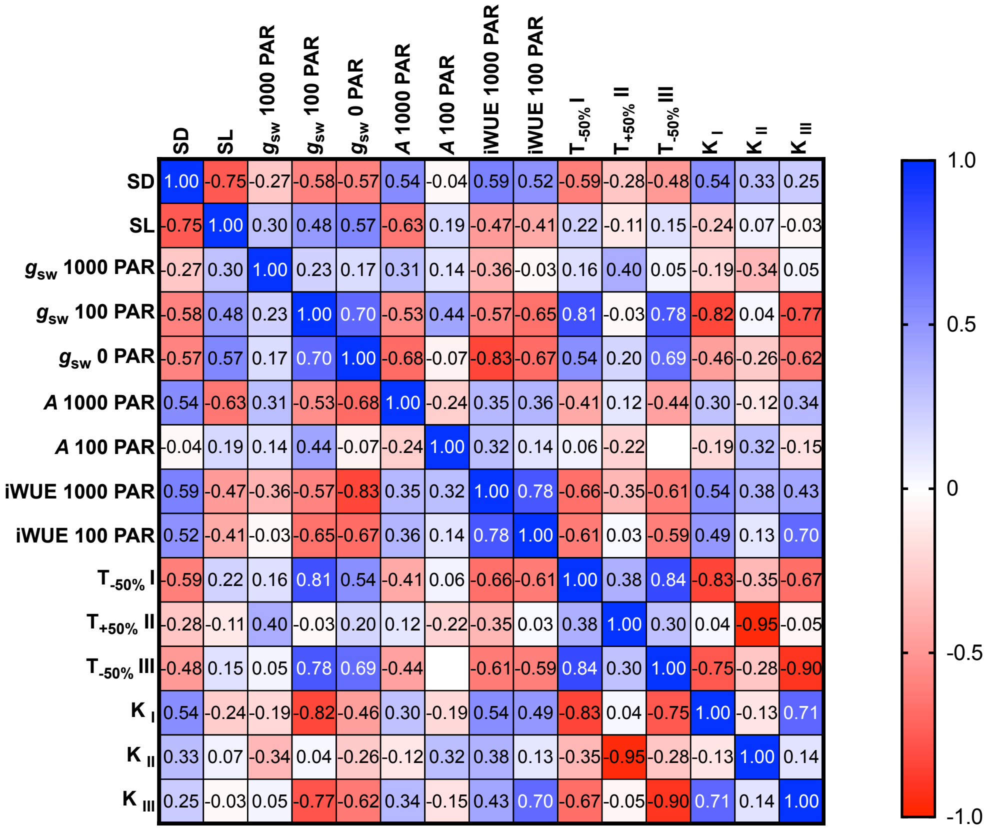


**Figure S4 - Anatomical influence on gas exchange and stomatal kinetics.** (A) Correlation matrix between anatomical traits (SD and SL), steady-state $g_{sw}$ (at 1000, 100 and 0 PAR), steady-state *A* (at 100 and 1000 PAR), iWUE (1000 and 100 PAR), half-time (for the light transitions 1000-100 ($T_{-50\%}$ I), 100-1000 ($T_{+50\%}$II) and 1000-0 PAR ($T_{-50\%}$III)) and constant rate (for the light transitions 1000-100 ($K_I$), 100-1000 ($K_{II}$) and 1000-0 PAR ($K_{III}$)); n=15.

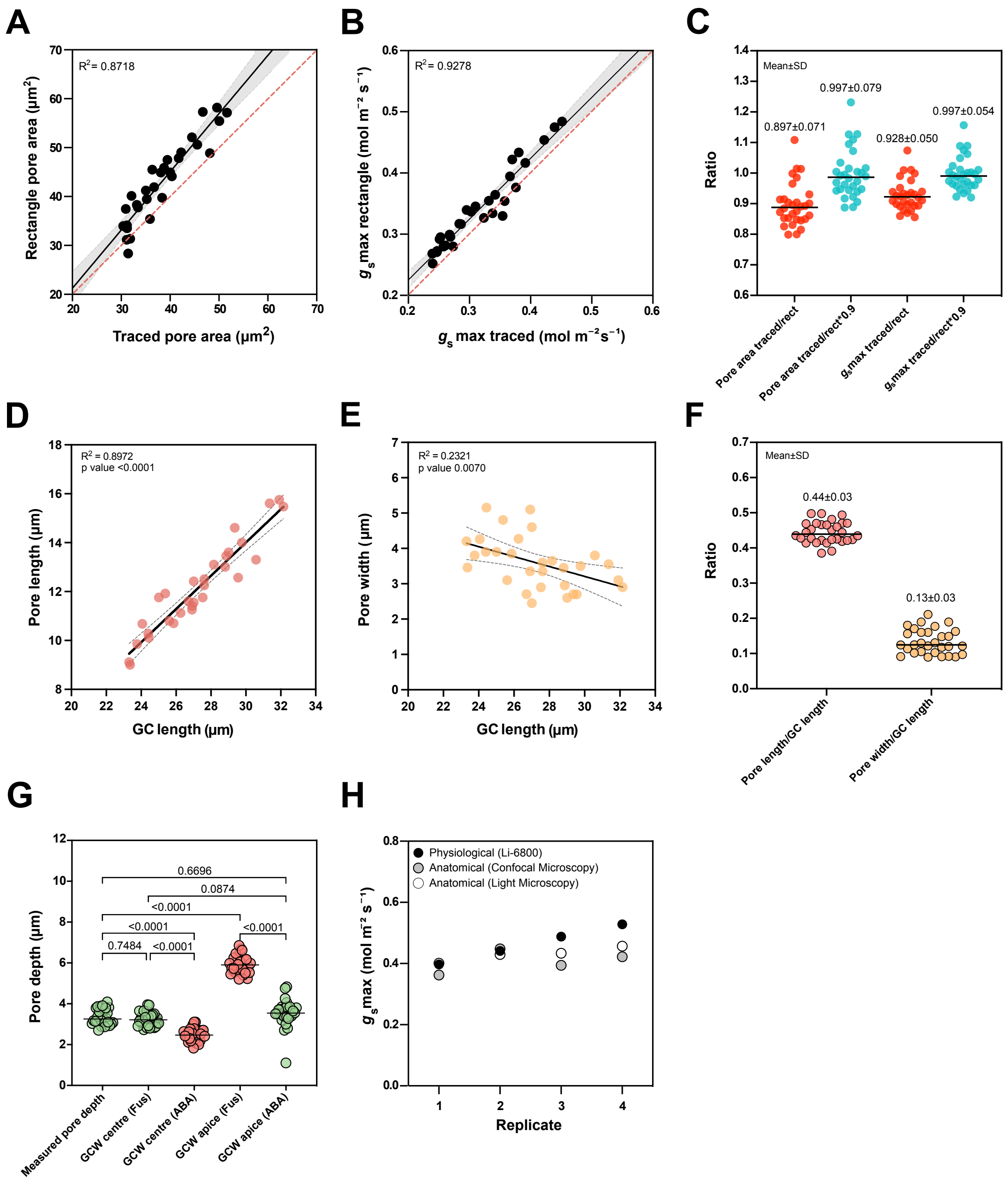


**Figure S5 - Quantitative morphometry of the graminoid stomata in *B. distachyon* Bd21-3.** (A) Linear regression between hand-traced pore area and rectangle pore area. (B) Linear regression of anatomical $g_s$max calculated with hand-traced pore area and with rectangle pore. (C) Ratios between hand-traced and rectangle pore, hand-traced and rectangle*0.9, anatomical gsmax calculated with hand-traced and rectangle pores, and anatomical gsmax calculated with hand-traced and rectangle*0.9 pores. (D) Linear regression between guard cell length (GCL) and pore length (PL). (E) Linear regression between guard cell length (GCL) and pore width (PW). (F) Ratios between GCL and PL and between GCL and PW. (G) Comparison of measured pore depth (l) with itself, $GCW_C$ (fusicoccin-treatment), $GCW_C$ (ABA-treatment), $GCW_A$ (fusicoccin-treatment) and $GCW_A$ (ABA-treatment). Green groups indicate that the measured parameters were a good indication of pore depth (l), red groups were not. (H) Physiological (IRGA-based), anatomical (light microscopy-based) and anatomical (confocal-based) $g_s$max values of each replicate shown in Fig. 5I. $R^2$ and p-values are indicated.

**Supplementary Dataset 1 - Gas exchange and anatomical data measured and analyzed in this study.**
[link]
